# Bioactive exometabolites drive maintenance competition in simple bacterial communities

**DOI:** 10.1101/2021.09.05.459016

**Authors:** John L. Chodkowski, Ashley Shade

## Abstract

During prolonged resource limitation, bacterial cells can persist in metabolically active states of non-growth. These maintenance periods, such as those experienced in stationary phase, can include upregulation of secondary metabolism and release of exometabolites into the local environment. As resource limitation is common of many environmental microbial habitats, we hypothesized that neighboring bacterial populations employ exometabolites to compete or cooperate during maintenance, and that these exometabolite-facilitated interactions can drive community outcomes. Here, we evaluated the consequences of exometabolite interactions over stationary phase among three environmental strains: *Burkholderia thailandensis* E264, *Chromobacterium subtsugae*, and *Pseudomonas syringae* pv.*tomato* DC3000. We assembled them into synthetic communities that only permitted chemical interactions. We compared the responses (transcripts) and outputs (exometabolites) of each member with and without neighbors. We found that transcriptional dynamics were changed with different neighbors, and that some of these changes were coordinated between members. The dominant competitor *B. thailandensis* consistently upregulated biosynthetic gene clusters to produce bioactive exometabolites for both exploitative and interference competition. These results demonstrate that competition strategies during maintenance can contribute to community-level outcomes. It also suggests that the traditional concept of defining competitiveness by growth outcomes may be narrow, and that maintenance competition could be an additional or alternative measure.

## Introduction

Bacteria interact with other bacteria and their environment within complex, multi-species communities. Bacterial interactions rely on the ability to sense and respond to both biotic and abiotic stimuli [1, 2]. These stimuli include physical, chemical or molecular cues, and can alter bacterial behaviors [3, 4], and ultimately, can also alter community functioning [5, 6]. It is expected that interspecies interactions play an important role in shaping microbial community dynamics [7]. However, multiple stimuli in the environment make it difficult to disentangle the separate influences of abiotic versus biotic stimuli on microbial community dynamics [8]. Therefore, efforts to characterize and distinguish community responses to biotic stimuli, such as those that facilitate interspecies interactions, will provide insights into the specific roles that microbial interactions play in shaping their communities [9].

Interspecies interactions can be facilitated through small molecules [10]. Extracellular small molecules are collectively referred to as exometabolites [11, 12, 13]. Depending on the exometabolite produced, these molecules can mediate interspecies interactions that range from competitive to cooperative [14]. Of these interaction types, competition has been shown to have a major influence in structuring microbial communities [15, 16, 17]. Thus, competitive interactions that are mediated by exometabolites are also expected to influence to microbial community dynamics. In addition, different types of exometabolites can be employed by bacteria to gain advantage in both exploitative (e.g. nutrient scavenging) and interference (direct cell damage) categories of competition.

Traditionally, competition has been viewed through the lens of resource acquisition [18]. In previous studies, competitiveness is modeled with respect to yield given resource consumption and growth [19, 20]. However, competition for *survival* or *maintenance* may be just as important as competition for yield, especially during periods of resource limitation [21, 22]. Competition during maintenance is likely common in some free-living environments, such as soils, sequencing batch reactors, and the human gut that experience long periods of nutrient famine punctuated by short periods of nutrient influx [23, 24, 25, 26]. The stationary phase of a bacterial growth curve falls within this context of growth cessation, and pulses of nutrients may be transiently available as cells die and lyse (necromass), while the total population size remains stagnant. Stationary phase is often coordinated with a metabolic shift to secondary metabolism [27, 28]. Therefore, an effective “maintenance” competitor may produce bioactive exometabolites, like antibiotics, which are often produced as a result of secondary metabolism. Bacteria can activate biosynthetic gene clusters (BSGCs) to produce bioactive exometabolites [29]. The activation of BSGCs is closely tied to stress responses, suggesting that bacteria can sense the stress of competition [30, 31]. While it is known that certain exometabolites can trigger BSGC upregulation and, more generally alter transcription [32], there is much to understand about the outcomes of interspecies interactions for BSGCs in multi-member microbial communities.

Here, we build on our previous research to understand how exometabolite-mediated interactions among bacterial neighbors contribute to community outcomes in a simple, three-member community (Table 1). These three members are commonly associated with terrestrial environments (soils or plants) and were chosen because of reported [33] and observed interspecies exometabolite interactions in the laboratory. We used a synthetic community (“SynCom”) approach [34] by applying our previously described transwell system [35], which allowed for evaluation of “community goods” within a media reservoir that was shared among members. The members’ populations were physically isolated by membrane filters at the bottom of each transwell, but could interact chemically via the reservoir. In our prior work, we investigated each member’s exometabolites and transcription over stationary phase, and the objective was to understand monoculture responses (in minimal glucose media) before assembling the more complex 2- and 3- member communities. We found that each member in monoculture produced a variety of exometabolites in stationary phase, including bioactive molecules involved in competition [36]. In this work, we build to 2- and 3- community memberships to ask: How do members interact via exometabolites in simple communities during maintenance (stationary phase), and what are the competitive strategies and outcomes of those interactions? What genetic pathways, molecules, and members drive the responses?

**Table 1.** Bacterial members used in the synthetic community (SynCom) system.

| Member | *Burkholderia thailandensis* E264 | *Chromobacterium subtsugae* ATCC 31532<sup>a</sup> | *Pseudomonas syringae* pv. *tomato* DC3000 |
| --- | --- | --- | --- |
| **Genome size (Mb)** | 6.72 | 4.76 | 6.54 |
| **Family** | *Burkholderiaceae* | *Neisseriaceae* | *Pseudomonadaceae* |
| **No. of CDSs<sup>b</sup>** | 5 639 | 4 393 | 5 576 |
| **Chromosomes** | 2 | 1 | 1 |
| **Plasmids** | 0 | 0 | 2 |
| **ATCC** | 700388 | 31532 | BAA-871 |
| **Reference** | 37 | 38 | 39 |

<sup>a</sup>This member in previous work [35, 36] was referred to as *Chromobacterium violaceum*.
This member has been reclassified [40].
<sup>b</sup>CDSs, Coding sequences.

We found that *B. thailandensis* had a major influence on the transcriptional responses of both *C. subtsugae* and *P. syringae*, and that this influence could be attributed to an increase in both interference and exploitative competition strategies. Furthermore, we observed non-additive transcriptional responses and exometabolite production, particularly for *B. thailandensis* with respect to BSGCs. These findings show that diverse competitive strategies can be deployed even when bacterial neighbors are surviving rather than exponentially growing. Therefore, we suggest that contact-independent, exometabolite-mediated interference and exploitation are important competitive strategies in resource-limited environments and support the non-yield outcome of maintenance.

## Materials and Methods

### Bacterial strains and culture conditions

We selected three environmental bacterial strains for the SynCom experiments that were originally isolated from various plant/soil habitats and that had prior evidence of exometabolite interactions among them in the laboratory [Table 1; 33, 37-40]. Freezer stocks of *B. thailandensis*, *C. subtsugae*, and *P. syringae* were plated on half-concentration Trypticase soy agar (TSA50) at 27°C for at least 24 h. Members were inoculated in 7 ml of M9–0.2% glucose medium and grown for 16 h at 27°C, 200 rpm. Cultures were then diluted into 50 ml M9-0.2% glucose medium such that exponential growth phase was achieved after 10 h of incubation at 27°C, 200 rpm. Members were diluted in 50 ml M9 glucose medium to target ODs (*B. thailandensis* 0.3 OD, *C. subtsugae*: 0.035 OD, *P. syringae* 0.035 OD). The high initial OD for *B. thailandensis* was necessary such that stationary phase would be achieved by all members within a 2 h window after 24 h incubation in the transwell plate. The glucose concentration in the final dilution varied upon community membership- 0.067% for monocultures, 0.13% for pairwise cocultures, and 0.2% for the 3-member community. For each member, 48 ml of diluted culture was transferred as 4 mL aliquots in 12, 5 mL Falcon tubes to more efficiently prepare replicate transwell plates.

### Synthetic community experiments

Transwell plate preparation was performed as previously described [35]. Briefly, we used sterile filter plates with 0.22-μm-pore polyvinylidene difluoride (PVDF) filter bottoms (Millipore MAGVS2210). Prior to use, filter plates were washed three times with sterile water using a vacuum apparatus (NucleoVac 96 vacuum manifold; Clontech Laboratories). The filter of well H12 was removed with a sterile pipette tip and tweezer, and 31 ml of M9 glucose medium was added to the reservoir through well H12. The glucose concentration in the reservoir varied upon community membership- 0.067% for monocultures, 0.13% for pairwise cocultures, and 0.2% for the 3-member community. Glucose concentration was adjusted to plate occupancy (e.g., 3-member communities had higher number of wells occupied than 2- or 1-member). Our aim was for each member to achieve stationary phase at similar times across all conditions to compare transcripts and exometabolites under similar growth trajectories. In other words, available resources were standardized while keeping the well occupancy for each member constant. With this design, transcripts and exometabolites in cocultures that deviated from those in monocultures could be attributed to interspecies interactions and not complicated by offset in member growth trajectories across the experimental conditions.

Each well was filled with 130 μL of culture or medium (prepared as described, above; see methods section: Bacterial strains and culture conditions). For each plate, a custom R script (RandomArray.R [see script at https://github.com/ShadeLab/PAPER_Chodkowski_mSystems_2017/blob/master/R_analysis/RandomArray.R]) was used to randomize community member placement in the wells so that each member occupied a total of 31 wells per plate. In total, there were 7 community conditions- 3 monocultures, 3 pairwise cocultures, and the 3-member community. Each member occupied 31 wells per plate regardless of experimental condition. Thus, “baseline” exometabolites could be determined in the monocultures, and then deviations in exometabolite abundance or detection in the cocultures could be attributed to interspecies interactions. A time course was performed for each replicate. The time course included an exponential phase time point (12.5 h) and 5 time points assessed every 5 h over stationary phase (25 h – 45 h). Four biological replicates were performed for each community condition for a total of 28 experiments. For each experiment, 6 replicate filter plates were prepared for destructive sampling for a total of 168 transwell plates.

Filter plates were incubated at 27°C with gentle shaking (∼0.32 rcf). For each plate, a custom R script (RandomArray.R [see script at https://github.com/ShadeLab/PAPER_Chodkowski_mSystems_2017/blob/master/R_analysis/RandomArray.R]) was used to randomize wells for each organism assigned to RNA extraction (16 wells) and flow cytometry (5 wells). The following procedure was performed for each organism when a transwell plate was destructively sampled: i) wells containing spent culture assigned to RNA extraction were pooled (∼100 μL/well) into a 1.7 mL microcentrifuge tube and flash frozen in liquid nitrogen and stored at −80 until further processing. ii) 20 μL from wells assigned for flow cytometry were diluted into 180 μL Tris-buffered saline (TBS; 20 mM Tris, 0.8% NaCl [pH 7.4]). In community memberships where *P. syringae* was arrayed with *B. thailandensis*, *P. syringae* had a final dilution of 70-fold in TBS. In community memberships where *P. syringae* was arrayed in monoculture or in coculture with *C. subtsugae*, *P. syringae* had a final dilution of 900-fold in TBS. Final dilutions for *B. thailandensis* and *C. subtsugae* were 1 300-fold and 1 540-fold, respectively. Each member was diluted differently to achieve a suitable events/second range on the flow cytometer for accurate cell counting. Populations were then stained and analyzed on the flow cytometer for live/dead counting (see Supplementary Methods). iii) Spent medium (∼31 ml) from the shared reservoir was transferred to 50 mL conical tubes, flash-frozen in liquid nitrogen and stored at −80 °C prior to metabolite extraction.

### RNA-seq

RNA extraction, sequencing, quality control, and count matrix generation was performed as previously published [36, see Supplementary Methods].

### Transcriptomics

#### Quality filtering and differential gene expression analysis

Count matrices for each member were quality filtered in two steps: genes containing 0 counts in all samples were removed, and genes with a transcript count of ≤10 in more than 90% of samples were removed. DESeq2 [41] was used to extract size factor and dispersion estimates. These estimates were used as external input into ImpulseDE2 for the analysis of differentially regulated genes [42]. ImpulseDE2 determines differential expression by comparing longitudinal count datasets. Case-control (Cocultures-monoculture control) analyses were analyzed to identify genes with differences in temporal regulation at an FDR-corrected threshold of 0.01. Genes that passed the FDR threshold were further filtered for genes that had at least one timepoint with a log2 fold-change (LFC) >= 1 or <= −1. Thus, we defined differentially expressed genes (DEGs) as genes that met both the FDR-corrected and LFC thresholds. For each member, differences in gene regulation between the three coculture conditions was visualized with Venn diagrams using the VennDiagram package [43].

Differentially expressed genes were first determined by comparing each coculture condition to the monoculture control and applying a LFC threshold (see above). We then determined a second set of DEGs by comparing pairwise cocultures to each other. ImpulseDE2 case-control analyses were performed as follows: *B. thailandensis* coculture with *C. subtsugae* (case) compared to *B. thailandensis* coculture with *P. syringae* (control), *C. subtsugae* coculture with *B. thailandensis* (case) compared to *C. subtsugae* coculture with *P. syringae* (control), and *P. syringae* coculture with *B. thailandensis* (case) compared to *P. syringae* coculture with *C, subtsugae* (control). Genes that passed the FDR-corrected threshold of 0.01 based on ImpulseDE2 analysis and had at least one time point with a LFC of >= 1 or <= −1 represented coculture specific DEGs. The DEGs determined from monoculture comparisons and coculture comparisons were then categorically grouped using Clusters of Orthologous Groups (COG).

#### COG analysis

Protein fasta files were downloaded from NCBI and uploaded to eggNOG-mapper v2 (http://eggnog-mapper.embl.de/) to obtain COGs. The DEGs determined from ImpulseDE2 and LFC thresholds were categorized as upregulated or downregulated based on temporal expression patterns. DEGs with consistent positive LFC throughout all stationary phase time points were categorized as upregulated. DEGs with consistent negative LFC throughout all stationary phase time points were categorized as downregulated. These DEGs were then assigned to COGs, grouped based on temporal up/downregulation patterns, and plotted using ggplot2 [44].

#### Principal coordinates analysis and statistics

We extracted DEGs based on our previously described definition. A variance-stabilizing transformation was performed on normalized gene matrices using the rlog function in DESeq2. A distance matrix based on the Bray-Curtis dissimilarity metric was then calculated on the variance-stabilized gene matrices and principal coordinates analysis was performed using the R package vegan [45]. Principal coordinates were plotted using ggplot2. Coordinates of the first two PCoA axes were used to perform PROTEST analysis using the PROTEST function in vegan. Dissimilarity matrices were used to perform PERMANOVA and variation partitioning and using the adonis and varpart functions in vegan, respectively. The RVAideMemoire package [46] was used to perform a post-hoc pairwise PERMANOVAs.

#### Biosynthetic gene cluster (BSGC) analysis

NCBI accession numbers were uploaded to antiSMASH 6 beta bacterial version [47] to identify genes involved in BSGCs using default parameters. Where possible, literature-based evidence and BSGCs uploaded to MIBiG [48] were used to better inform antiSMASH predictions. Log2 fold-changes (LFCs) were calculated for all predicted biosynthetic genes within each predicted cluster by comparing coculture expression to monoculture expression at each time point. Average LFCs were calculated from all predicted biosynthetic genes within a predicted BSGC at each time point. Temporal LFC trends were plotted using ggplot2. An upregulated BSGC was defined as a BSGC that had at least two consecutive time points in stationary phase with a LFC > 1.

#### Network analysis

Unweighted co-expression networks were created from quality filtered and normalized expression data. Networks were generated for pairwise cocultures containing *B. thailandensis*. First, data were quality filtered as previously described (see methods section: *Quality filtering and differential gene expression analysis*). Then, normalized expression data was extracted from DESeq2. Twenty-three and twenty-four RNA-seq samples from each member were used for network analysis in the *B. thailandensis*-*C. subtsugae and B. thailandensis*-*P.syringae* cocultures, respectively (24 samples/member; 6 timepoints, 4 biological replicates). Only 23 samples were used in the *B. thailandensis*-*C. subtsugae* network analysis because RNA-seq failed for *C. subtsugae* at 45 h, biological replicate 2. Interspecies networks were then inferred from the expression data using the context likelihood of relatedness [49] algorithm within the R package Minet [50]. Gene matrices for each coculture pair were concatenated to perform the following analysis. Briefly, the mutual information coefficient was determined for each gene-pair. To ensure robust detection of co-expressed genes, a resampling approach was used as previously described [51]. Then, a Z-score was computed on the mutual information matrix. A Z-score threshold of 4.5 was used to determine an edge in the interspecies network. Interspecies networks were uploaded into Cytoscape version 3.7.1. for visualization, topological analysis, and enrichment analysis [52].

Gene annotation and gene ontology (GO) files were obtained for *B. thailandensis*, *P. syringae,* and *C. subtsugae* for enrichment analyses. For *B. thailandensis,* annotation and ontology files were downloaded from the Burkholderia Genome Database (https://www.burkholderia.com). For *P. syringae*, annotation and ontology files were downloaded from the Pseudomonas Genome Database (http://www.pseudomonas.com/strain/download). Annotation and ontology files for *C. subtsugae* were generated using Blast2GO version 5.2.5 [53]. InterProScan [54] with default parameters were used to complement gene annotations from *C. subtsugae*. GO terms were assigned using Blast2GO with default parameters. In addition, genes involved in secondary metabolism were manually curated and added to these files as individual GO terms. These genes were also used to update the GO term GO:0017000 (antibiotic biosynthetic process), composed of a collection of all the biosynthetic genes. (see methods section: *Biosynthetic gene cluster (BSGC) analysi*s).

Topological analysis was performed as follows: Nodes were filtered from each coculture network to only select genes from one member. The GLay community cluster function in Cytoscape was used to determine intra-member modules. Functional enrichment analysis was then performed on the modules using the BiNGO package [55] in Cytoscape.

To determine interspecies co-regulation patterns, we filtered network nodes that contained an interspecies edge. Functional enrichment analysis was performed on the collection of genes containing interspecies edges for each member using the BiNGO package in Cytoscape. Genes contained within modules of interest (e.g. modules containing either thailandamide or malleilactone genes) were selected in Cytoscape for visualization. The biosynthetic gene cluster organization of thailandamide and malleilactone were obtained from MIBig and drawn in InkScape.

Protein sequences from an interspecies gene of interest (CLV_2968) within a network module that also contained thailandamide genes from the *B. thailandensis*-*C. subtsugae* network and an interspecies gene of interest (PSPTO_1206) within a network module that also contained malleilactone genes from the *B. thailandensis*-*P. syringae* network were obtained. A protein blast for each protein was run against *B. thailandensis* protein sequences. *B. thailandensis* locus tags were extracted from the top blast hit from each run. Normalized transcript counts for these 4 genes of interest were plotted in R. Time course gene trajectories were determined using a loess smoothing function.

### Metabolomics

#### LCMS, feature detection, and quality control

Standard operating protocols were performed at the Department of Energy Joint Genome Institute as previously described [36]. MZmine2 [56] was used for feature detection and peak area integration as previously described [36]. Select exometabolites were identified in MZmine2 by manual observation of both MS and MS/MS data. We extracted quantities of these identified exometabolites for ANOVA and Tukey HSD post-hoc analysis in R. We filtered features in three steps to identify coculture-accumulated exometabolites. The feature-filtering steps were performed as follows on a per-member basis: (i) retain features where the maximum peak area abundance occurred in any of the coculture communities ; (ii) a noise filter, the minimum peak area of a feature from a replicate at any time point needed to be 3 times the maximum peak area of the same feature in one of the external control replicates, was applied; (iii) coefficient of variation (CV) values for each feature calculated between replicates at each time point needed to be less than 20% across the time series.

Four final feature data sets from polar and nonpolar analyses in both ionization modes were analyzed in MetaboAnalyst 5.0 [57], as reported in our prior work [36, see Supplementary Methods].

#### Principal coordinates analysis and statistics

A distance matrix based on the Bray-Curtis dissimilarity metric was used to calculate dissimilarities between exometabolite profiles. Principal coordinates analysis was performed using the R package vegan. Principal coordinates were plotted using ggplot2. Coordinates of the first two PCoA axes were used to perform Protest analysis using the protest function in vegan. Dissimilarity matrices were used to perform PERMANOVA and variation partitioning and using the adonis and varpart functions in vegan, respectively. The RVAideMemoire package was used to perform a post-hoc pairwise PERMANOVAs. Monoculture controls were removed to focus on coculture trends.

## Results

### Overview

Our major data types included both transcriptomics and metabolomics, and we integrate these to interpret SynCom dynamics and interactions. Our longitudinal design resulted in 288 RNAseq samples across the three members, and 168 community metabolomics samples analyzed in each of four mass spectral modes (polar/nonpolar, positive/negative modes = 672 total mass spectral profiles). After quality control, we were left with 281 RNAseq and 605 total mass spectral profiles for the integrated analyses [https://github.com/ShadeLab/Paper_Chodkowski_3member_SynCom_2021/tree/master/SummaryOfSamples]. First, we present results of general responses of transcription (section 1) and exometabolomics (section 2), separately. Then, we integrate transcriptomic and metabolomic efforts to determine the upregulation of biosynthetic gene clusters (BSGCs) and identify exometabolites of interest from mass spectrometry (section 3). We then present a transcriptomics co-expression network to ask how the upregulation of BSGCs influenced interspecies interactions through coordinated longitudinal gene expression (section 4). Lastly, we discuss non-additive outcomes in both transcriptional responses and exometabolite output (section 5).

#### 1. Stationary phase transcript dynamics of microbial community members

We had four replicate, independent timeseries of each of seven community memberships (three monocultures plus three cocultures of every pair and the 3-member community), and here focus on the coculture analyses to gain insights into community outcomes (Figs. S1 & S2). Differential expressed genes were determined by comparing time series transcript trajectories applying an FDR and LFC threshold (see methods: *Quality filtering and differential gene expression analysis*). First, we compared each coculture to the monoculture control. A range of 153 to 276 genes were differentially expressed by each member in coculture, irrespective of the identity of neighbors (Fig. S3). In addition, each member also had differential gene expression that was unique to a particular neighbor(s). Summarizing across all cocultures, 1089/5639 (19.3%), coding sequences (CDSs), 1991/4393 CDSs (45.3%), and 3274/5576 CDSs (58.7%) DEGs were determined for *B. thailandensis*, *C. subtsugae*, and *P. syringae*, respectively. Both community membership and time contributed to the transcriptional response of each member (Fig. 1, Table S1). Together, these data suggest that there are both general and specific consequences of neighbors for the transcriptional responses of these bacterial community members.

**Figure 1.**
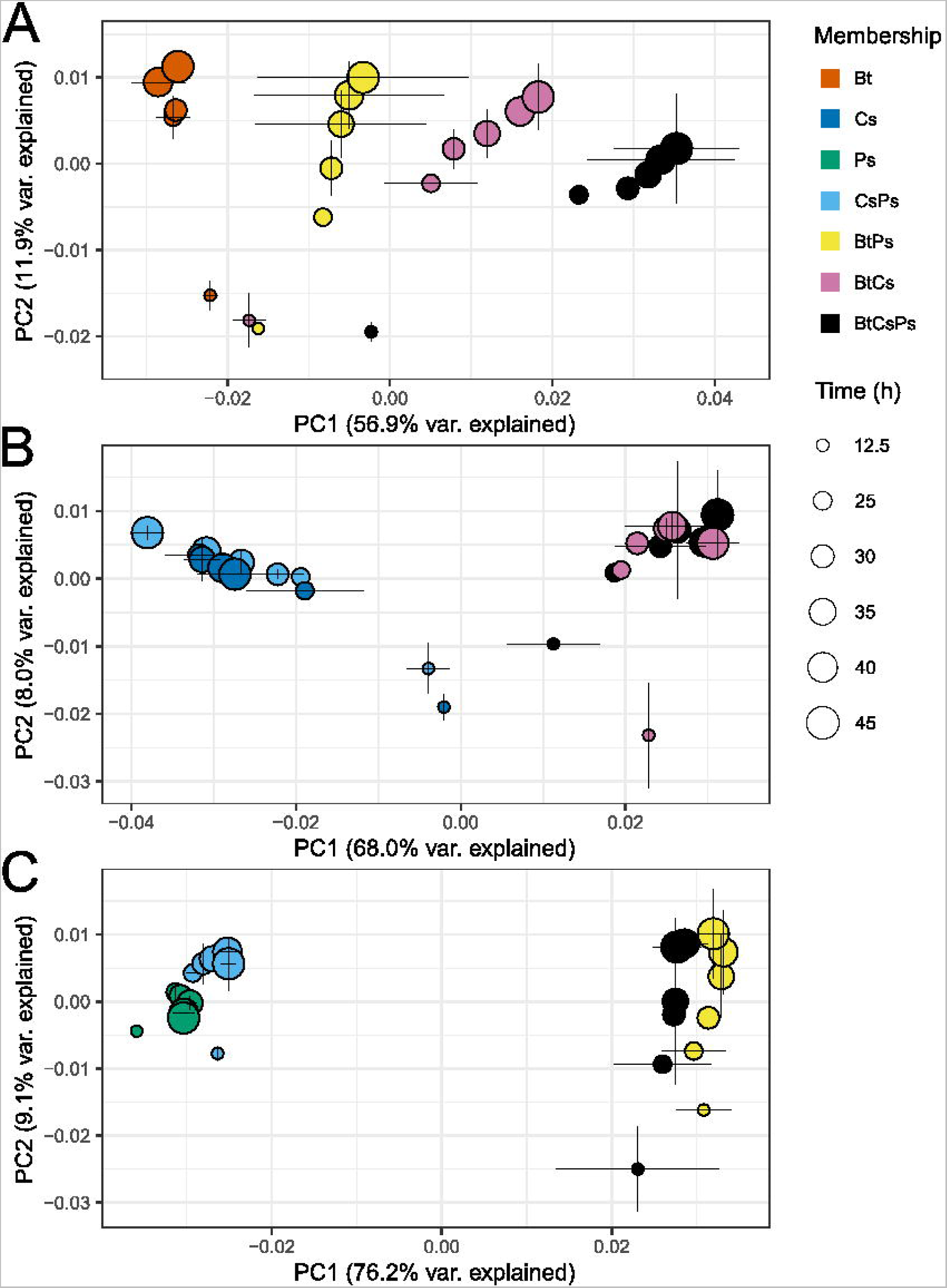
Transcriptional responses are driven by community membership and time. Shown are principal coordinates analysis (PCoA) plots for *B. thailandensis* (Bt, A), *C. subtsugae* (Cs, B), and *P. syringae* (Ps, C). Each point represents a mean transcript profile for a community member given a particular community membership (neighbor(s) included, indicated by symbol color) and sampled at a given time point over exponential and stationary phases (in hours since inoculation, h, indicated by symbol size, n=3 to 4 replicates per timepoint/community membership). The Bray-Curtis distance metric was used to calculate dissimilarities between transcript profiles. Error bars are 1 standard deviation around the mean axis scores.

Temporal trajectories in transcript profiles were generally reproducible across replicates for each member given a particular community membership (PROTEST analyses, Table S2). Each member had a distinct transcript profile (0.480L≤Lr2 ≤ 0.778 by Adonis; P value, 0.001; all pairwise false discovery rate [FDR]-adjusted P values, ≤0.01 except for two community memberships, Table S3). For all ordinations, community membership had the most explanatory value (Axis 1), followed by time (Axis 2), with the most variation explained by the interaction between time and membership (Table S1). Membership alone accounted for 60.6% and 77.0% of the variation explained in *C. subtsugae* and *P. syringae* analyses, respectively and 46.3% in the *B. thailandensis* analysis (Table S1).

When included in the community, *B. thailandensis* strongly determined the transcript profiles of the other two members. For example, the inclusion of *B. thailandensis* in a coculture differentiated transcript profiles for both *C. subtsugae* and *P. syringae* (Fig. 1B & 1C). Thus, *B. thailandensis* appears to have had a dominating influence on the transcriptional response of neighbors, and these responses were dynamic with respect to time.

We analyzed clusters of orthologous groups of proteins (COGs) to infer the responses of members to their neighbors. Differentially expressed genes were categorized as upregulated or downregulated based on temporal patterns and representation in COGs (Fig. S4). We focused on the largest differences between total DEGs upregulated and total DEGs downregulated within a COG, which provides insights into broad biological processes affected by community membership. COGs with large differences toward upregulation in *B. thailandensis* included cell motility [N], secondary metabolites biosynthesis, transport, and catabolism [Q], and signal transduction mechanisms [T] while COGs with large differences toward downregulation included defense mechanisms [V], energy production and conversion [C], translation, and ribosomal structure and biogenesis [J]. These results suggest that *B. thailandensis* responds to neighbors via downregulation of growth and reproduction and upregulation of secondary metabolism. We therefore hypothesized that *B. thailandensis* was producing bioactive exometabolites against *C. subtsugae* and *P. syringae* to competitively inhibit their growth.

Because of the strong transcript response of *C. subtsugae* and *P. syringae* when neighbored with *B. thailandensis* (Fig. 1B & 1C), we focused on COGs within community memberships with *B. thailandensis* (Fig. S4B & S4C, rows 2 & 3). The COG with large differences toward upregulation in both *C. subtsugae* and *P. syringae* were translation, ribosomal structure and biogenesis [J]. COG groups tending toward downregulation in *C. subtsugae* and *P. syringae* were signal transduction mechanisms [T] and secondary metabolites biosynthesis, transport, and catabolism [Q], respectively. These results suggest that the presence of *B. thailandensis* alters its neighbor’s ability to respond to the environment and inhibits secondary metabolism.

The above analyses focused on DEGs determined by comparing each coculture to the monoculture control. However, we also wanted to understand differences between pairwise-coculture memberships to determine if the alterations in transcripts were attributed to specific memberships (aka interspecies interactions). A total of 436, 1 762, and 2 962 DEGs were determined when comparing the *B. thailandensis* pairwise-cocultures, the *C. subtsugae* pairwise-cocultures, and the *P. syringae* pairwise-cocultures, respectively. Categorizing coculture DEGs based COGs and temporal up/downregulation gene trends shows membership-specific effects (Fig. S5). These data suggest coculture-specific transcriptional alterations driven by interspecies interactions. Given the physical separation of members in our SynCom plate system, these interspecies interactions were very likely exometabolite-mediated.

#### 2. Stationary phase exometabolite dynamics of microbial communities

Because member populations are physically separated in the SynCom transwell system but allowed to interact chemically, observed transcript responses in different community memberships are inferred to result from exometabolite interactions. Spent medium from the shared medium reservoir was collected from each transwell plate and analyzed using mass spectrometry to detect exometabolites. Our previous manuscript focused on exometabolite dynamics in monocultures [36]. Here, we focused our analysis on those exometabolites that had maximum accumulation in a coculture (either in pairs or in 3-member community). Consistent with the transcript analysis, we found that both community membership and time explained the exometabolite dynamics, and that the explanatory value of membership and time was maintained across all polarities and ionization modes (Fig. 2, Table S4).

**Figure 2.**
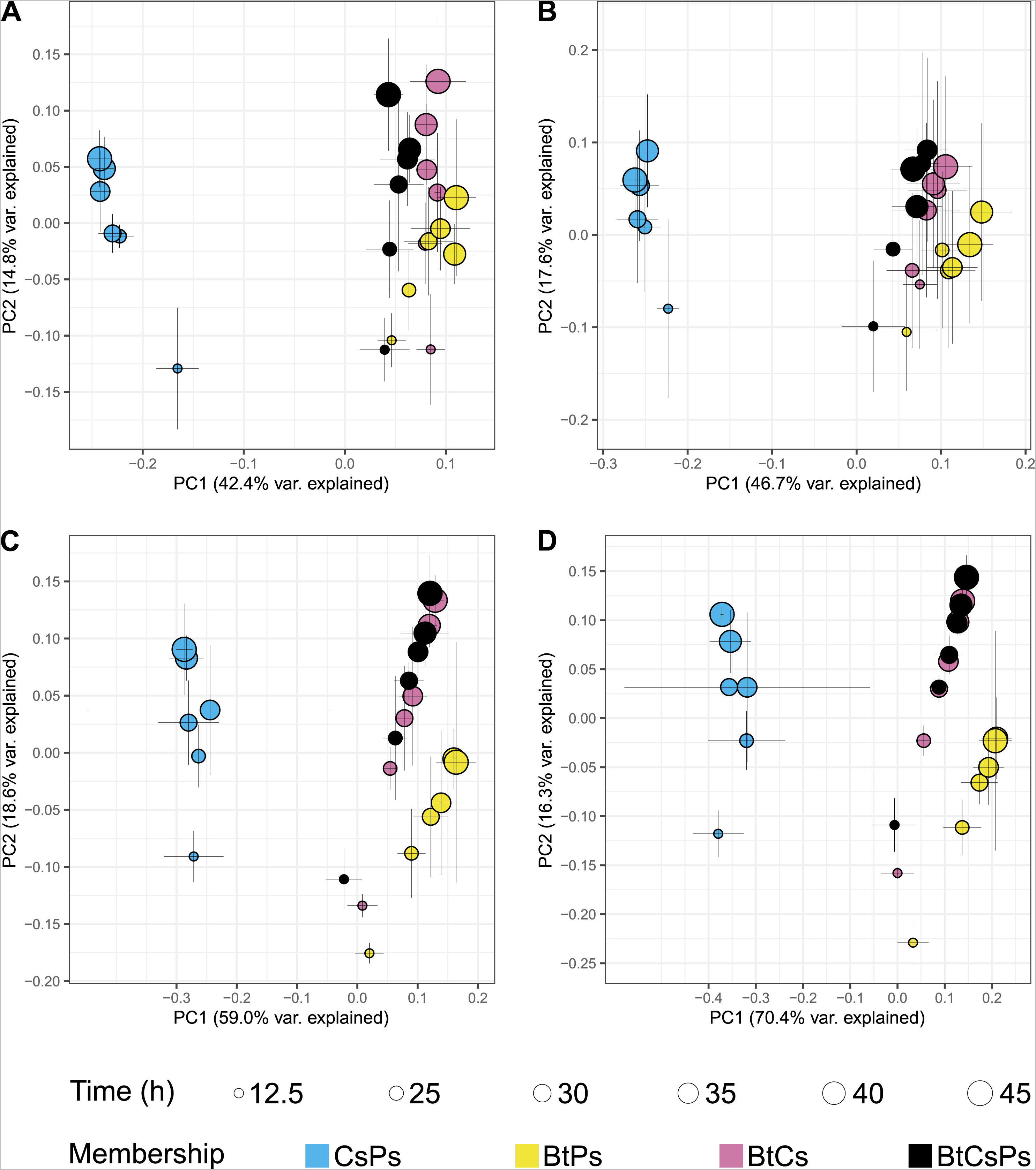
Bacterial community exometabolite profiles differ by community membership and time. Shown are PCoA plots for exometabolite profiles from the following mass spectrometry modes: polar positive (A), polar negative (B), nonpolar positive (C), and nonpolar negative (D). Each point represents the mean exometabolite profile (relative contributions by peak area) given a particular community membership (indicated by symbol color) at a particular time point (indicated by symbol shape). The Bray-Curtis distance metric was used to calculate dissimilarities between exometabolite profiles. Error bars are 1 standard deviation around the mean axis scores (n= 2 to 4 replicates). Bt is *B. thailandensis*, Cs is *C. subtsugae*, and Ps is *P. syringae*.

Temporal trajectories in exometabolite profiles were generally reproducible across replicates with some exceptions (PROTEST analyses, Table S5, Supplementary File 1). Exometabolite profiles were distinct by community membership (0.475L≤Lr2 ≤ 0.662 by Adonis; P value, 0.001; all pairwise false discovery rate [FDR]-adjusted P values, ≤0.01 except for two comparisons, Table S6), and also dynamic over time. As observed for the member transcript profiles, and the interaction between membership and time had the highest explanatory value for the exometabolite data (Table S4).

We found that the *C. subtsugae*-*P. syringae* coculture exometabolite profiles were consistently the most distinct from the other coculture memberships (Fig. 2), supporting, again, that the inclusion of *B. thailandensis* was a major driver of exometabolite dynamics, possibly because it provided the largest or most distinctive contributions to the community exometabolite pool. Indeed, we observed that a majority of the most abundant exometabolites were either detected uniquely in the *B. thailandensis* monoculture or accumulated substantially in its included community memberships (Fig. 3). Some exometabolites detected in *B. thailandensis-*inclusive communities were not detected in its monocultures (Fig. 3D), suggesting that the inclusion of neighbors contributed to the accumulation of these particular exometabolites (e.g. upregulation of biosynthetic gene clusters or lysis products). *C. subtsugae* and *P. syringae* contributed less to the 3-member community exometabolite profile, as exometabolites detected in the *C. subtsugae*-*P. syringae* coculture were less abundant and had lower accumulation over time in the 3-member community (Fig. 3A). Together, these results suggest that *B. thailandensis* can suppress or overwhelm expected outputs from neighbors.

**Figure 3.**
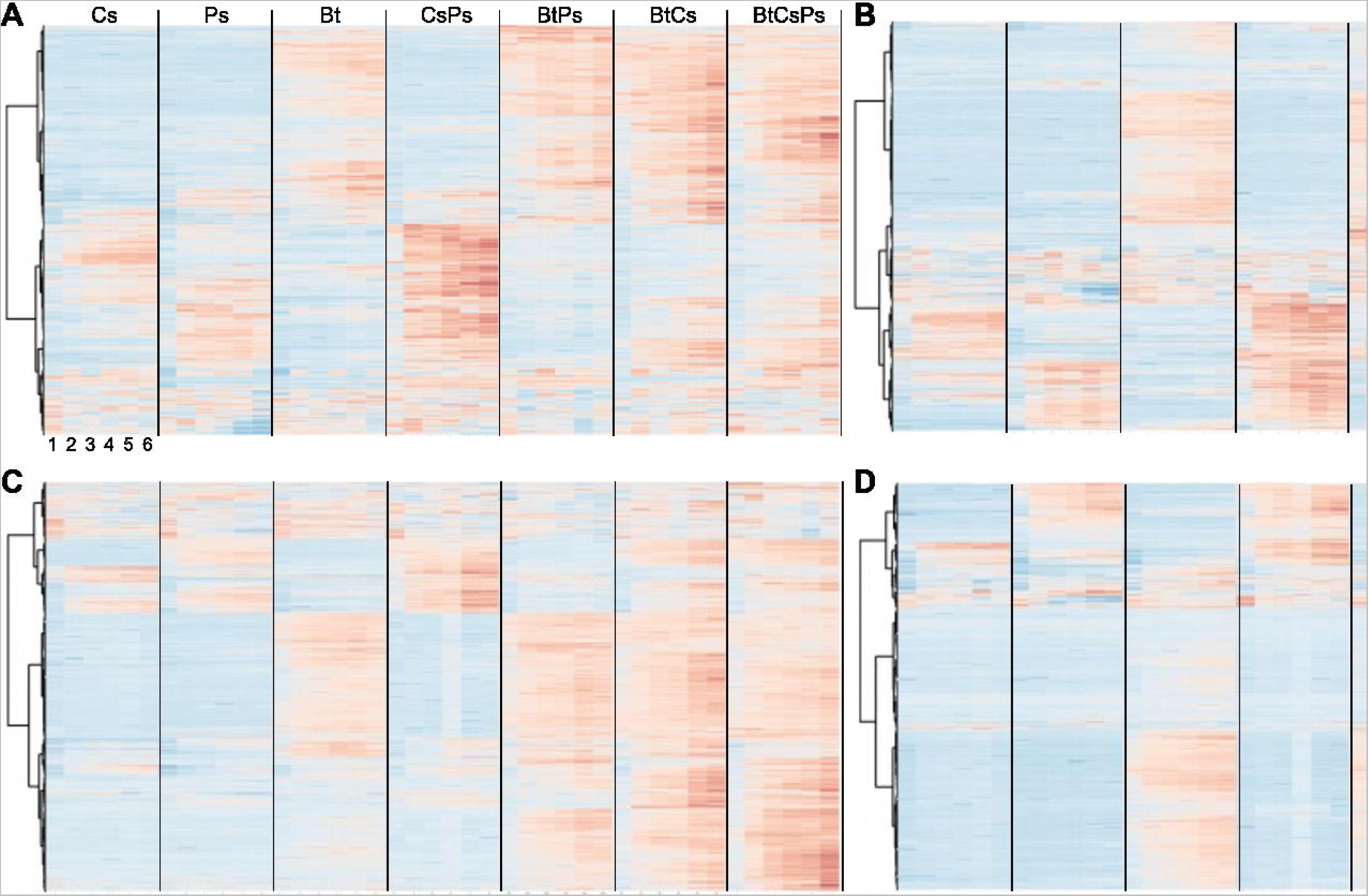
Exometabolites have membership-specific production and temporal accumulation. A heat map of coculture accumlated exometabolites is shown for polar positive (A), polar negative (B), nonpolar positive (C), and nonpolar negative (D) modes, for *C. subtsugae* monoculture (Cs), *P. syringae* monoculture (Ps), *B. thailandensis* monoculture (Bt), *C. subtsugae-P. syringae* coculture (CsPs)*, B. thailandensis*-*P. syringae* coculture (BtPs)*, B. thailandensis*-*C. subtsugae* coculture (BtCs), and the 3- member community (BtCsPs), where samples are in columns and exometabolites are in rows. Data for each sample are the averages from independent time point replicates (nL=L2 to 4).

In summary, we observed both increased accumulation and unique production of exometabolites in pairwise cocultures and in the 3-member community, with *B. thailandensis* contributing the most to the shared exometabolite pool as determined by comparisons with its monoculture exometabolite profile. Related, the transcriptional responses of *C. subtsugae* and *P. syringae* in the 3-member community is most similar to their respective transcriptional response when neighbored with *B. thailandensis* alone, despite the presence of the third neighbor.

#### 3. B. thailandensis increases competition strategies in the presence of neighbors

We observed relatively unchanged viability in *B. thailandensis* (Fig. S6). On the contrary, we observed a slight reduction (∼2.1 log2 fold change) in *C. subtsugae* live cell counts, and a drastic reduction (∼4.7 log2 fold change) in *P. syringae* live cell counts, when either member was cocultured with *B. thailandensis* (Figs. S7 & S8; monoculture cell counts vs *B. thailandensis* coculture cells counts). We note that one doubling occurred in *B. thailandensis* and *P. syringae* monocultures, and in *C. violaceum* in pairwise coculture with *P. syringae.* We elaborate on this finding as the possibility of a reductive cell division as described in our previous manuscript [36]. Given this reduction in viability and that there have been competitive interactions between *B. thailandensis* and *C. subtsugae* previously reported [33], we hypothesized that *B thailandensis* was using competition strategies to influence its neighbors via production of bioactive exometabolites. If true, we would expect transcriptional upregulation in *B. thailandensis* biosynthetic gene clusters (BSGC) that encode bioactive exometabolites. Indeed, we found evidence of this when *B. thailandensis* had neighbors (Fig. 4, Table S7). This suggests that *B. thailandensis* responded to neighbors by upregulating genes involved in the production of bioactive compounds, likely to gain a competitive advantage. However, not all BSGCs in *B. thailandensis* were upregulated. Some BSGCs were unaltered or downregulated (Fig. S9). *C. subtsugae* upregulated only 1 BSGC in coculture with *B. thailandensis,* while *P. syringae* did not upregulate any BSGC in any coculture (Figs. S10 & S11). Interestingly, coculturing with *C. subtsugae* and *P. syringae* resulted in the upregulation of an unidentified beta-lactone and an unidentified non-ribosomal peptide synthetase (NRPS) in *B. thailandensis*, respectively. Similarly, coculturing with *B. thailandensis* resulted in the upregulation on an unidentified NRPS-Type I polyketide synthase in *C. subtsugae*. We also note that two additional unidentified NRPS passed the LFC threshold of 1 in *C. subtsugae*. However, these were only upregulated at the exponential phase time point and subsequently downregulated or below the LFC threshold in all stationary phase time points. Interspecies interactions led to the upregulation of BSGC in both *B. thailandensis* and *C. subtsugae* and three of these BSGC encode potentially novel bioactive exometabolites.

**Figure 4.**
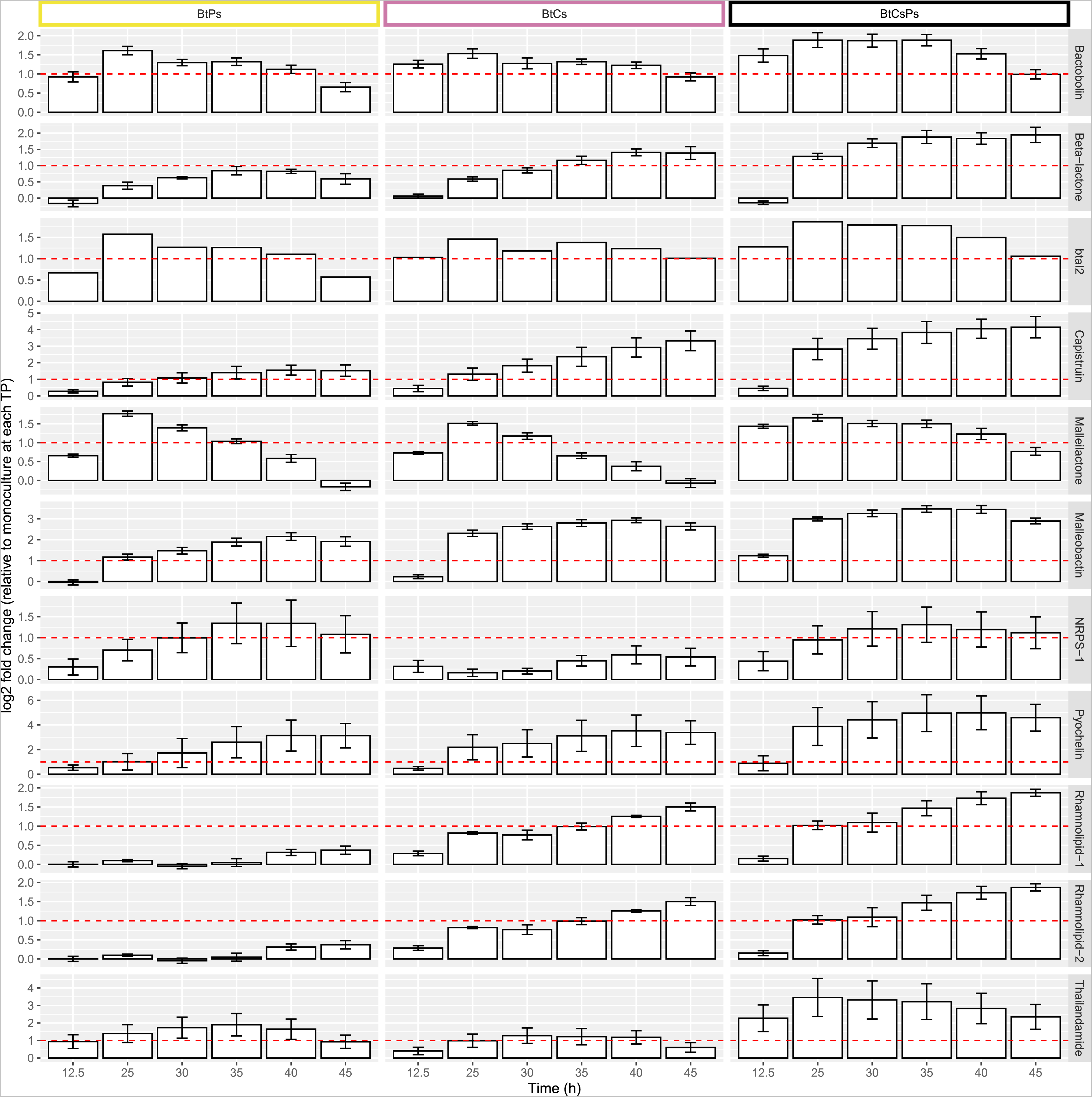
*B. thailandensis* upregulates biosynthetic gene clusters (BSGC) in cocultures. Columns represent community membership where Bt: *B. thailandensis*, Cs: *C. subtsugae*, and Ps: *P. syringae* and rows represent BSCC in *B. thailandensis*. Genes part of a BSGC were curated from antiSMASH predictions and literature-based evidence. Within each BSGC, log2 fold-changes (LFC) were calculated by comparing gene counts from a coculture to the monoculture control at each time point. LFC were then averaged from all biosynthetic genes in the BSGC at each time point. We defined an upregulated BSGC as a BSGC that had at least two consecutive stationary phase time points with a LFC > 1 (indicated by the horizontal line). Note that plots for each BSGC have separate scales for the Y-axis.

We were able to identify 6 of the 11 exometabolites from the upregulated *B. thailandensis* BSGCs and quantify their abundances from mass spectrometry data (Fig. 5, Supplementary File 2). For any given identified exometabolite, it differentially accumulated between community memberships containing *B. thailandensis* (Table S8), particularly when comparing the *B. thailandensis* monoculture compared to each coculture (Table S9). As expected, these identified exometabolites were not detected in community memberships that did not include *B. thailandensis*. Bactobolin was the only identified exometabolite that accumulated in monoculture to equivalent levels of accumulation in all coculture conditions. Thus, *B. thailandensis* increased competition strategies with neighbors through the upregulation and production of many bioactive exometabolites. Of these bioactive exometabolites, three are documented antimicrobials [58, 59, 60], two are siderophores [61, 62], and one is a biosurfactant [63]. We conclude that *B. thailandensis* produced bioactive exometabolites to competitively interact using both interference and exploitative competition strategies [64]. Given that *B. thailandensis* upregulated competition strategies, and responded more broadly in producing competition-supportive exometabolites when grown with neighbors, we hypothesized that these bioactive exometabolites are responsible for the altered transcriptional responses in *C. subtsugae* and *P. syringae*.

**Figure 5.**
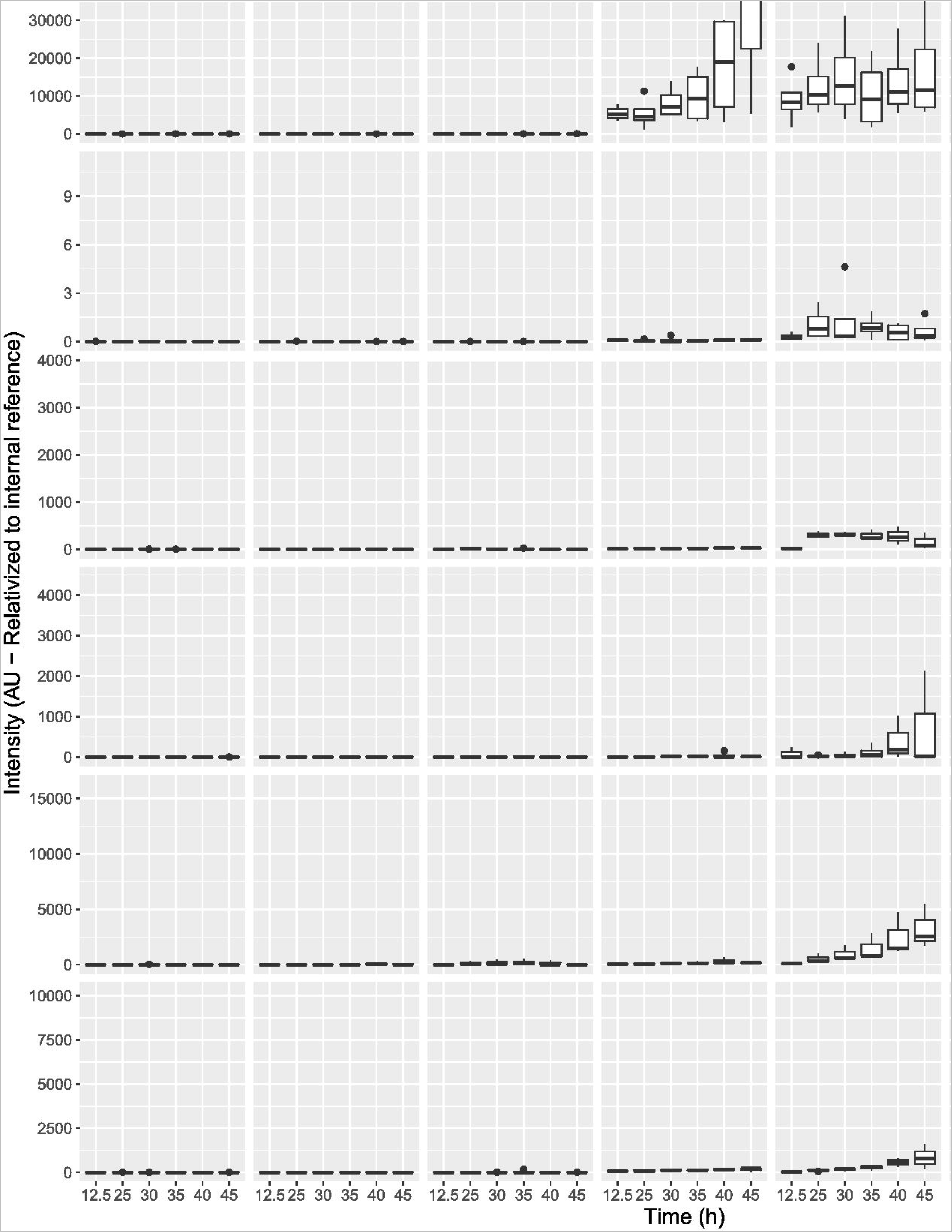
Identified exometabolites from *B. thailandensis* BSGC temporally accumulate in cocultures. Known bioactive secondary metabolites produced by *B. thailandensis* were identified in mzMine2 through the observation of MS and MS/MS data. The accumulation of each exometabolite was quantified through time (n = 2-4 integrated peak areas per time point). The bottom and top of the box are the first and third quartiles, respectively, and the line inside the box is the median. The whiskers extend from their respective hinges to the largest value (top), and smallest value (bottom) was no further away than 1.5× the interquartile range.

In our experimental design, we adjusted glucose concentration depending on plate occupancy. Glucose concentration increased as plate occupancy increased (31 wells vs 62 wells vs 93 wells), but a member consistently occupied 31 wells across all experimental conditions. One complication of this design is that population density and resource concentration could contribute to differences in transcripts and exometabolites in a member-agnostic manner. To address this, we performed additional SynCom experiments to affirm confidence that some changes in transcripts and exometabolites are attributable to exometabolite-mediated interspecies interactions. In these experiments, we increased the plate occupancy of *B. thailandensis* in monoculture while subsequently increasing resource concentration. Pairwise cocultures and the 3-member community SynCom experiments were repeated as well (see Supplementary methods). We calculated the relative gene expression of three genes in the thailandamide operon (*thaF*, *thaK*, and *thaQ*) through RT-qPCR by comparing each experimental condition to the monoculture control (*B. thailandensis*, 31 wells in M9-0.067% glucose). Decreased gene expression was observed across all three genes as both plate occupancy and resource concentration increased in *B. thailandensis* monocultures. In fact, *thaF*, *thaK*, and *thaQ* gene expression was further reduced in the 93 well *B. thailandensis* monoculture compared to the 62 well *B. thailandensis* monoculture, suggesting that the thailandamide operon trended towards reduced expression as a function of *B. thailandensis* plate occupancy in monoculture conditions. On the contrary, *thaF*, *thaK*, and *thaQ* had increased expression in all coculture memberships, suggesting that exometabolite interspecies interactions were responsible for the increased expression of a BSGC in *B. thailandensis* (Table S10).

#### 4. Interspecies co-transcriptional networks reveal coordinated gene expression related to competition

We performed interspecies co-expression network analysis to infer interspecies interactions. We used temporal profiles from transcriptional responses to generate co-expression networks for *B. thailandensis*-*C. subtsugae* and *B. thailandensis*-*P. syringae* cocultures, respectively (Table S11). As expected, the majority of nodes network had intraspecies edges only, with interspecies edges comprising 1.85% and 1.90% of the total edges in the *B. thailandensis*-*C. subtsugae* and *B. thailandensis*-*P. syringae* networks, respectively. We explored interspecies edges for evidence of interspecies transcriptional co-regulation.

We performed two analyses (module analysis and GO enrichment) to validate networks and infer interspecies interactions (Fig. S12). Module analysis validated networks as intraspecies modules enriched for biological processes (Supplementary File 3). To infer interspecies interactions, we filtered genes with interspecies edges and performed enrichment analysis (Supplementary File 4). The top enriched Gene Ontology (GO) term for *B. thailandensis* when paired with *C. subtsugae* was antibiotic synthesis of thailandamide, supporting interference competition. Though the top enriched GO term in *B. thailandensis* when paired with *P. syringae* was bacterial-type flagellum-dependent cell motility, antibiotic synthesis of malleilactone was also enriched. Both thailandamide genes from the *B. thailandensis*-*C. subtsugae* network (Fig. 6) and malleilactone genes from the *B. thailandensis*-*P. syringae* network (Fig. S13) formed near-complete modules within their respective BSGCs.

**Figure 6.**
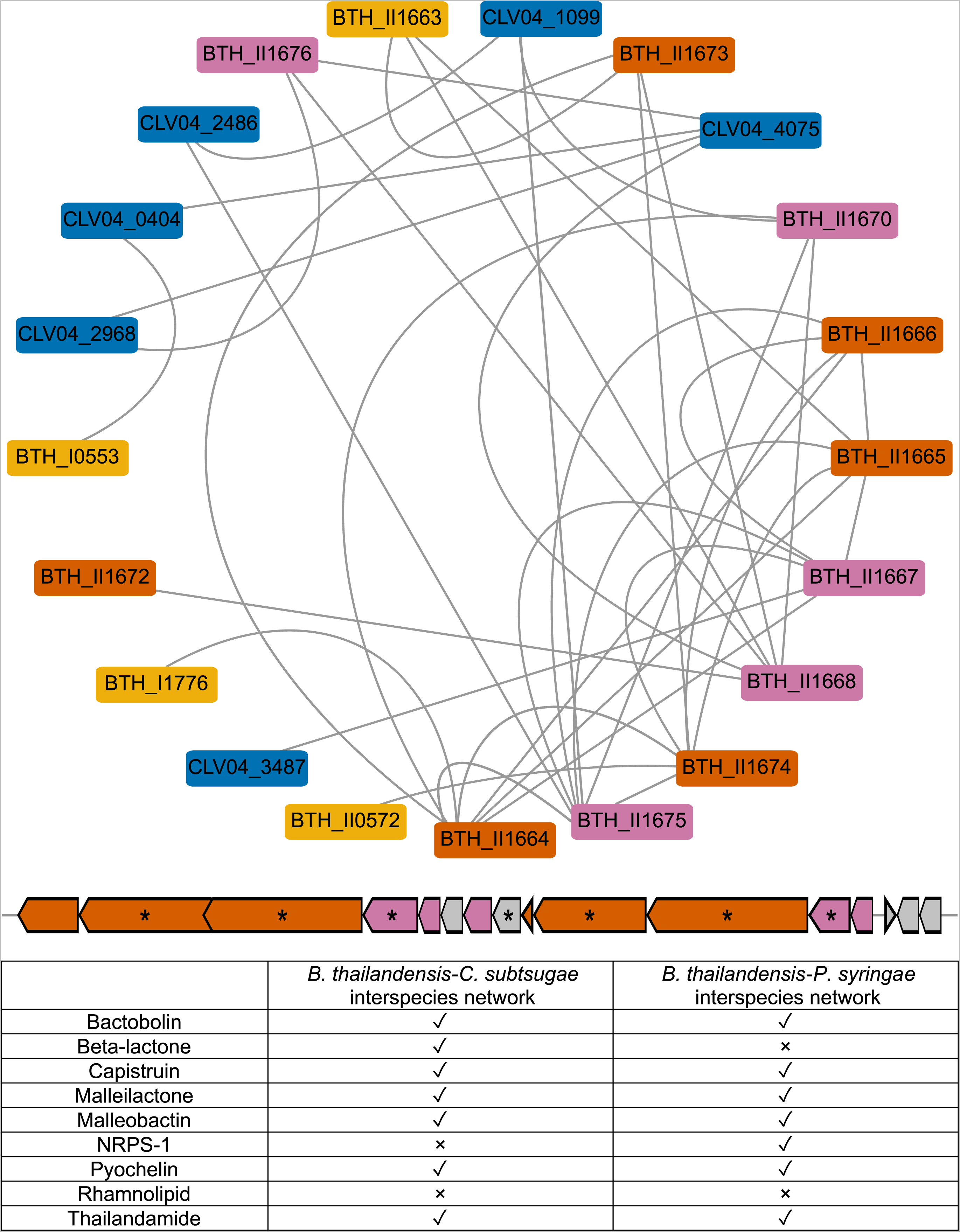
*B. thailandensis* genes involved in thailandamide production are detected as interspecies edges in the *B. thailandensis*-*C. subtsugae* co-expression network and biosynthetic genes organize into network modules. A network module containing the thailandamide BSGC is shown. The network module nodes are color coded by *B. thailandensis* gene type (BSGC or not) and type of connections (interspecies or not): thailandamide biosynthetic genes that had interspecies edges (magenta), thailandamide biosynthetic genes that did not have interspecies edges (orange), or other genes that were not part of the BSGC (yellow); as well as genes that were from *C. subtsugae* (blue). The chromosomal organization of the thailandamide BSGC is shown below the network module. The same colors are applied to the BSGC operon. The operons also depict genes that were not detected within the interspecies network, shown in gray. Asterisks indicate core biosynthetic genes in the BSGCs, as predicted from antiSMASH. The table shows upregulated *B. thailandensis* BSGCs (Fig. 4) and whether there were interspecies edges detected (check is yes, x is no).

At least one gene from each of *B. thailandensis’s* upregulated BSGCs (Fig. 4) had an interspecies edge, except for rhamnolipid. The top GO term for both *C. subtsugae* and *P. syringae* genes that had edges shared *B. thailandensis* was bacterial-type flagellum-dependent motility. Other notable enriched GO processes were efflux activity for *C. subtsugae* and signal transduction for *P. syringae*. Specifically, a DNA starvation/stationary phase gene (CLV04_2968, Fig. 6), *dspA*, was within the network module that also contained thailandamide genes from the *B. thailandensis*-*C. subtsugae* network and a TonB-dependent siderophore receptor gene (PSPTO_1206, Fig. S13) was within the network module that also contained malleilactone genes from the *B. thailandensis*-*P. syringae* network (Supplementary File 5). Interestingly, both CLV04_2968 and PSPTO_1206 were DEGs and downregulated when cocultured with *B. thailandensis* (Figs. S14A & S15A, respectively). Additionally, the closest homolog for *dspA* in *B. thailandensis* was unaltered (BTH_I1284, Supplementary File 6) when cocultured with *C. subtsugae* (Fig. S14B) and the closest homolog to the TonB-dependent receptor in *B. thailandensis* (BTH_I2415, Supplementary File 7) was a DEG and upregulated when cocultured with *P. syrinage* (Fig. S15B). Taken together, these co-expression networks revealed interspecies coordinated expression patterns.

## Discussion

Here, we used a synthetic community system to understand how exometabolomic interactions determine members transcriptional responses and exometabolite outputs. Our experiment used a bottom-up approach to compare the seven possible community memberships of three members, and their dynamics in member transcripts and community exometabolites over stationary phase. Differential gene expression across community memberships and over time show that the exometabolites released by a member were sensed and responded to by its neighbors. Furthermore, members’ ouputs in monocultures changed because of coculturing, as evidenced by differential exometabolite production. The largest transcriptional alterations in *C. subtsugae* and *P. syringae* occurred when cocultured with *B. thailandensis*. Global expression patterns in *C. subtsugae* and *P. syringae* when in the 3-member community still resembled expression patterns in pairwise cocultures with *B. thailandensis.* These transcriptional alterations in *C. subtsugae* and *P. syringae* were coordinated with increases in *B. thailandensis* competitive strategies (evaluated by BSGC transcript upregulation and exometabolite abundance) . Numerous transcripts and exometabolite were non-additive in the 3-member community, suggesting that predictions of members outcomes in more complex communities will not always be a simple summation of pairwise outcomes. That interactions within a relatively simple community altered the transcriptional responses and exometabolite outputs of each member is important because these kinds of alterations could, in turn, drive changes in community structure and/or function in an environmental setting. For example, it was shown that interspecies interactions more strongly influenced the assembly of *C. elegans* gut communities than host-associated factors [65]. Therefore, mechanistic and ecological characterization of interspecies interactions will inform as to the principles that govern emergent properties of microbial communities.

Overall, competitive interactions predominated in this synthetic community. Our previous study found that, over stationary phase in monocultures, each member released and accumulated at least one exometabolite documented to be involved in either interference or exploitative competition [36]. This suggests that entry into stationary phase primed members for competitive interactions, whether there were heterospecific neighbors present. We interpret this strategy of preemptive aggression to be especially advantageous to *B. thailandensis,* as it successfully used competitive strategies against both *C. subtsugae* and *P. syringae*. *B. thailandensis’s* success was supported by decreased viable *P. syringae* cells when cocultured with *B. thailandensis*. Though *C. subtsugae* viable cell counts were not as affected directly by the coculture with *B. thailandensis*, *B. thailandensis*-produced bactobolin [66] was detected in the shared medium reservoir. Bactobolin is a bacteriostatic antibiotic previously shown to be bioactive against *C. subtsugae* [33] through ribosome binding [58]. But, *C. subtsugae* can resist bactobolin through upregulation of an RND-type efflux pump [67]. This finding also is supported by our data, as all genes coding for the CdeAB-OprM RND-type efflux system were DEGs and upregulated in *C. subtsugae* cocultures with *B. thailandensis* (CLV04_2413-CLV04_2415).

When cocultured with *B. thailandensis*, we observed COG groups such as translation, ribosomal structure and biogenesis [J] had large differences toward upregulation in both *C. subtsugae* and *P. syringae*. At first glance, this seems at odds with our interpretation of *B. thailandensis* competitiveness toward *C. subtsugae* and *P. syringae*. In other words, how is *B. thailandensis* effectively competing via interference competition if both *C. subtsugae* and *P. syringae* are upregulating machinery for growth? There is both theoretical [68] and experimental [69] evidence that show how cells treated with antibiotics stimulate ribosomal production to maintain a sufficient number of active ribosomes. As previously mentioned, *B. thailandensis*-produced bactobolin binds to the ribosome and can inhibit *C. subtsugae* [33, 58]. We also have evidence that bactobolin inhibits *P. syringae* (data not shown). It could be that bactobolin is stimulating ribosomal production in *C. subtsugae* and *P. syringae* as a survival mechanism to maintain protein production by maintaining enough active ribosomes.

Coculturing can induce secondary metabolism [70, 71, 72] because an exometabolite produced by one microbe can prompt secondary metabolism in a neighbor [31]. We found that coculturing led to the upregulation of numerous BSGCs in *B. thailandensis*. These exometabolites included bactobolin, malleilactone [61, 73; siderophore and cytotoxin], malleobactin [74, 75; siderophore], capistruin [76; lasso peptide], thailandamide [77; polyketide], pyochelin [62; siderophore], rhamnolipids [63; biosurfactants], and two uncharacterized BSGCs encoding nonribosomal peptide synthetases. Of these exometabolites, bactobolin, capistruin, and thailandamide have documented antimicrobial activities through translation inhibition [58], transcription inhibition [59], and inhibition of fatty acid synthesis [60], respectively. For those exometabolites we were able to identify with mass spectrometry, their accumulation in cocultures was correlated with the upregulation of their BSGCs. Furthermore, up/downregulated patterns across all *B. thailandensis* BSGCs is consistent with ScmR global regulatory patterns of secondary metabolism [78]. Though we were not able to pinpoint the exact inducers of BSGCs, exometabolites such as antibiotics [79] and primary metabolites [80] have been documented to induce secondary metabolism in *B. thailandensis*. *C. subtsugae* can inhibit *B. thailandensis* [33] but we did not observe *B. thailandensis* inhibition based on cell counts. However, we did find that in stationary phase *C. subtsugae*-*B. thailandensis* cocultures, *C. subtsugae* upregulated an uncharacterized hybrid nonribosomal peptide synthetase-type I polyketide synthase. *P. syringae* was the least competitive of the three neighbors, as evidenced by a reduction in live cell counts when cocultured with *B. thailandensis*. Also, *P. syringae* did not increase competitive strategies when cocultured, as no BSGCs were upregulated across all coculture conditions. In summary, though all three neighbors had potential to use competitive strategies and maintained competitive strategies in monoculture [36], *B. thailandensis* was most successful in cocultures over stationary phase through increased production of exometabolites involved in interference and exploitative competition strategies.

Given the upregulation of BSGCs in *B. thailandensis* and the strong transcriptional responses of *C. subtsugae* and *P. syringae* to the presence of *B. thailandensis*, we hypothesized that competitive exometabolites were contributing to their community dynamics. Thus, we used a co-expression network analysis with our longitudinal transcriptome series to infer interspecies interactions [81]. This use of this approach was first demonstrated to infer coregulation between a phototroph-heterotroph commensal pair [82]. Our network confirmed that *B. thailandensis* BSGCs had coordinated gene expression patterns with both *C. subtsugae* and *P. syringae*. Interspecies nodes in both networks contained various genes involved in the aforementioned upregulated *B. thailandensis* BSGCs. In particular, we focused on interspecies edges within thailandamide nodes for the *B. thailandensis*-*C. subtsugae* network and interspecies edges within malleilactone nodes for the *B. thailandensis*-*P. syringae* network because these were significantly enriched as interspecies nodes. A *C. subtsugae* gene of interest, CLV04_2968, was contained within the thailandamide cluster of interspecies nodes. This gene codes for a DNA starvation/stationary phase protection protein and had the highest homology to the Dps protein in *Escherichia coli* across all *C. subtsugae* protein coding genes. Dps mediates tolerance to multiple stressors and *dps* knockouts are more susceptible to thermal, oxidative, antibiotic, iron toxicity, osmotic, and starvation stressors [83]. Interestingly, CLV04_2968 was downregulated when cocultured with *B. thailandensis*, suggesting that *B. thailandensis* attenuates *C. subtsugae* stress tolerance over stationary phase. While we observed a slight decrease in viable *C. subtsugae* cells when cocultured with *B. thailandensis*, one may expect *C. subtsugae* to have increased sensitivity to a subsequent stress [e.g. pH stress; 84] resulting from CLV04_2968 downregulation in the presence of *B. thailandensis*.

In the *B. thailandensis*-*P. syringae* co-expression network, a *P. syringae* gene of interest, PSPTO_1206, was contained within the malleilactone cluster of interspecies nodes. PSPTO_1206 is annotated as a TonB-dependent siderophore receptor. A *P. syringae* iron-acquistion receptor had coordinated expression with a malleilactone, which has been characterized as a siderophore with antimicrobial properties [61]. Interestingly, this gene was downregulated when in coculture with *B. thailandensis*. In contrast, the closest TonB-dependent siderophore receptor homolog to PSPTO_1206 in *B. thailandensis*, BTH_I2415, was upregulated in coculture conditions with *P. syringae*. To summarize, co-expression network analysis revealed interspecies coordinated gene expression patterns. Though determining directionality was beyond the scope of this analysis, we observed *B. thailandensis*-increased competition strategies were coordinated with a potential decrease in competition strategies in *C. subtsugae* via reduced stress tolerance and in *P. syringae* with reduced iron acquisition ability.

One feature of our study is that we adjusted glucose concentration depending on plate occupancy. Glucose concentration increased as membership increased, but a member consistently occupied 31 wells across all experimental conditions. One could argue that resource concentration contributed to differences in transcripts and exometabolites and not interspecies interactions. However, DEGs were present when comparing pairwise coculture conditions and these were attributed to differences in temporal regulation of COG categories (Fig. S5). More specifically, regarding BSGCs, an unidentified NRPS was upregulated in *B. thailandensis* when cocultured with *P. syringae* but not when cocultured with *C. subtsugae* (Fig. 4) and, an unidentified NRPS-Type I polyketide synthase was upregulated in *C. subtsugae* when cocultured with *B. thailandensis* but not when cocultured with *P. syringae* (Fig. S10). These differences occurred in experimental conditions where the glucose concentration was the same. Furthermore, we performed additional SynCom experiments where we increased the plate occupancy of *B. thailandensis* in monoculture while subsequently increasing resource concentration. Decreased gene expression was observed across all three RT-qPCR tested thailandamide genes as both plate occupancy and resource concentration increased in *B. thailandensis* monocultures. These same three genes had increased gene expression across all cocultures. These findings show that some undefined exometabolite interspecies interactions were responsible for the increased expression of a BSGC in *B. thailandensis*. Previous work has pinpointed central metabolites and antibiotics as elicitors of secondary metabolism [79, 80] but these mechanistic associations were beyond this study. Overall, we acknowledge that resource concentration and exometabolite output are intertwined, and subsequent work could test how initial resource availability determines SynCom outcomes.

A major goal in microbial ecology is to predict community dynamics for purposes of modulating and/or maintaining ecosystem function [85, 86]. At its core, microbial functional properties emerge, in part, from the concerted interactions of multi-species assemblages. The SynCom system provides a tractable experimental system to understand the relationships between exometabolite interactions and environmental stimuli to inform higher-order community interactions. Higher-order interactions are those that are unexpected based on interactions observed in simpler situations (e.g., of member pairs) [87, 88, 89]. Therefore, integrating different system variables, like transcriptome and metabolome dynamics, within controlled microbial communities will inform how unexpected phenomena arise and how they contribute to deviations in predictive models of community outcomes.

Our results indicated that each member continued to maintain competitive strategies despite stagnant population growth. *B. thailandensis* upregulated various bioactive exometabolites involved in both interference and exploitative competition when with neighbors. An effective competitor is often defined as by its ability to outcompete neighbors via growth advantage that stems from efficient nutrient uptake and/or biomass conversion rates [90, 91]. We add to this that a competitor can also have a fitness advantage through effective maintenance, which can similarly employ interference or exploitative competitive strategies despite no net growth. Maintenance may ensure survival in some environments that impose a stationary phase lifestyle, where long periods of nutrient depletion are punctuated with short periods of nutrient flux. In these scenarios, it warrants to understand how competitive strategies are deployed in the interim of growth and the extent to which these interactions contribute to long-term community outcomes. Though population levels remain constant, sub-populations of growing cells have been observed in stationary phase [92], and continued production of competitive exometabolites may serve as an advantageous strategy to hinder growth of competitors. In addition, some antibiotics remain effective in non-replicating bacteria [93]. The ability for continued maintenance via effective competition strategies during stationary phase may provide spatiotemporal maintenance of population levels before growth resumption [94]. Alternatively, both growth and non-growth strategies may be occurring simultaneously (e.g. as can occur in biofilms). The heterogeneity of biofilms may provide an environment where a bacterial population contains both stationary cells in the center of the colony with growing cells at the periphery of the colony that compete and alter developmental patterns of neighboring populations [95, 96]. Thus, we expect that insights into the long-term consequences of competition for microbial community outcomes will be gained by considering competition in both active growth and maintenance scenarios.

## Code availability

Computing code, workflows, and data sets are available at [https://github.com/ShadeLab/Paper_Chodkowski_3member_SynCom_2021]. R packages used during computing analyses included DEseq2 [41], ImpulseDE2 [42], VennDiagram [43], ggplot2 [44], vegan 2.5-4 [45], RVAideMemoire [46], Minet [50], rtracklayer [97], viridis [98], and helper functions [99, 100, 101, 102].

## Data availability

Genomes for *B. thailandensis*, *C. subtsugae*, and *P. syringae* are available at the National Center for Biotechnology Information (NCBI) under accessionnumbers <u>NC_007651</u> (Chromosome I)/<u>NC_007650</u> (Chromosome II), <u>NZ_PKBZ01000001</u>, and <u>NC_004578</u> (Chromosome)/<u>NC_004633</u> (Plasmid A)/<u>NC_004632</u> (Plasmid B), respectively. An improved annotated draft genome of *C. subtsugae* is available under NCBI BioProject accession number <u>PRJNA402426</u> (GenBank accession number <u>PKBZ00000000</u>). Data for resequencing efforts for *B. thailandensis* and *P. syringae* are under NCBI BioProject accession numbers <u>PRJNA402425</u> and <u>PRJNA402424</u>, respectively. Metabolomics data and transcriptomics data are also available at the JGI Genome Portal [103] under JGI proposal identifier 502921. MZmine XML parameter files for all analyses can be viewed at and downloaded from GitHub (see Dataset 7 at https://github.com/ShadeLab/Paper_Chodkowski_MonocultureExometabolites_2020/t <u>ree/master/Datasets</u>). Large data files (e.g., MZmine project files) are available upon request. Supplementary files are also available on GitHub (https://github.com/ShadeLab/Paper_Chodkowski_3member_SynCom_2021/tree/master/Supplemental_Files).

## Acknowledgements

This material is based upon work supported by the National Science Foundation under grant DEB 1749544 and by Michigan State University. In addition, metabolite analysis and transcript sequencing were provided by a DOE-JGI Community Science Program award (proposal identifier 502921). The work conducted by the U.S. Department of Energy Joint Genome Institute, a DOE Office of Science User Facility, is supported under contract number DE-AC02-05CH11231. J.L.C. was supported by the Eleanor L. Gilmore Fellowship from the Department of Microbiology and Molecular Genetics.

We thank Katherine B. Louie and Benjamin P. Bowen for support in mass spectral analysis.

## Author contributions

J.L.C. and A.S. conceived of and designed the study. J.L.C. performed the research and analyses. J.L.C. and A.S. wrote the manuscript.

