## Supplementary Methods for "Bioactive exometabolites drive maintenance competition in simple bacterial communities"

Chodkowski JL and A Shade.

Code and data:

<https://github.com/ShadeLab/Paper_Chodkowski_3member_SynCom_2021>

**Supplementary Methods**

This supplementary information contains expanded Materials and Methods.

**RNA-seq**

*RNA extraction*

RNA was extracted using the E.Z.N.A. Bacterial RNA kit (Omega Bio-tek, Inc.). An in-tube DNase I (Ambion, Inc AM2222, 2U) digestion was performed to remove DNA from RNA samples. RNA samples were purified and concentrated using the Qiagen RNAeasy MinElute Clean up Kit (Qiagen, Inc). Ten random samples were chosen to assess RNA integrity (RIN > 7) on an Agilent 2100 Bioanalyzer. Standard operating protocols were performed at the Department of Energy Joint Genome Institute as previously described [1].

*RNA sample prep, sequencing, QC, read preprocessing, and filtering*

Standard operating protocols were performed at the Department of Energy Joint Genome Institute as previously described [1].

*Pseudoalignment and counting*

Reads from each library were pseudoaligned to the transcriptome of each member with kallisto [2]. Raw counts from each library were combined into a gene count matrix for each member. The gene count matrix was used for downstream analyses.

**Flow cytometry**

Diluted cultures were stained with the Thermo Scientific LIVE/DEAD BacLight bacterial viability kit at final concentrations of 1.5 μM Syto9 (live stain) and 2.5 μM propidium iodide (dead stain). Two hundred microliters of stained cultures were transferred to a 96-well microtiter U-bottom microplate (Thermo Scientific). Twenty microliters of sample were analyzed on a BD Accuri C6 flow cytometer (BD Biosciences) at a fluidics rate of 66 μl/min and a threshold of 500 on an FL2 gate. The instrument contained the following optical filters: FL1-533, 30 nm; FL2-585, 40 nm; and FL3, 670-nm longpass. The counting accuracy of the flow cytometer was checked with green fluorescent protein beads (Thermo Scientific). Data were analyzed using BD Accuri C6 software version 1.0.264.21 (BD Biosciences).

**Metabolomics**

*Normalization and heatmap analysis*

Features were normalized by an ITSD reference feature (see Dataset 5 at <https://github.com/ShadeLab/Paper_Chodkowski_MonocultureExometabolites_2020/tree/master/Datasets>) and cube root transformed. Reference features for polar analyses in positive ([^13^C,^15^N]proline) and negative ([^13^C,^15^N]alanine) modes were determined by the ITSD with the lowest CV value across all samples. The reference feature for nonpolar data sets was the ITSD ABMBA. Heat maps were generated in MetaboAnalyst using Ward’s clustering algorithm with Euclidean distances from Z-scored data. Data for each sample are the averages from independent time point replicates (n = 2 to 4). Normalized and transformed data sets were exported from MetaboAnalyst to generate principal-coordinate analysis (PCoA) plots in R.

**Effects of plate occupancy and resource concentration on gene expression**

*SynCom experiments*

Additional SynCom experiments (6 conditions, 3 replicates/condition), were prepared as described (see methods section: Bacterial strains and culture conditions and Synthetic Community Experiments). The conditions varied based on plate occupancy (# of wells occupied by each member) and resources (% glucose) in the transwell plate. The conditions were as follows: *B. thailandensis* (31 wells) in M9-0.067% glucose, *B. thailandensis* (62 wells) in M9-0.13% glucose, *B. thailandensis* (93 wells) in M9-0.2% glucose, *B. thailandensis-C. subtsugae* (31 wells/member) in M9-0.13% glucose, *B. thailandensis- P. syringae* (31 wells/member) in M9-0.13% glucose, and *B. thailandensis-C. subtsugae- P. syringae* (31 wells/member) in M9-0.2% glucose. Plates were destructed after 45 h incubation and the following procedures were performed: 1) Wells containing spent culture from each member were separately pooled into 15 mL conical tubes, flash frozen in liquid nitrogen, and stored at -80 until further processing. 2) Spent medium (~31 ml) from the shared reservoir was transferred to 50 mL conical tubes, flash-frozen in liquid nitrogen and stored at −80 °C.

*RNA extraction, QC, and cDNA synthesis*

RNA was extracted using the E.Z.N.A. Bacterial RNA kit (Omega Bio-tek, Inc.). An in-tube DNase I (Ambion, Inc. AM2222, 2U) digestion was performed to remove DNA from RNA samples. RNA samples were purified and concentrated using the Qiagen RNAeasy MinElute Clean up Kit (Qiagen, Inc.). RNA samples were quantified on a Qubit using the RNA High Sensitivity Assay Kit (Thermo Fisher Scientific, Inc.). RNA samples were then sent to the RTSF Genomics Core at Michigan State University for high sensitivity RNA ScreenTape analysis on an Agilent 4200 TapeStation. TapeStation analysis confirmed successful digestion of DNA. Total RNA (150 ng/sample) was synthesized to cDNA using the Invitrogen SuperScript III First-Strand Synthesis kit (Thermo Fisher Scientific, Inc.). cDNA samples were quantified by Qubit in preparation of target genes for RT-qPCR.

Three genes from the *B. thailandensis* thailandamide operon were targeted for relative quantification normalized to the *rpoD* reference gene. Primers used for RT-qPCR are shown in Table S12. We first confirmed amplification of intended targets. Each of these genes were amplified from *B. thailandensis* gDNA (100 ng) using the Phusion High-Fidelity DNA Polymerase (New England Biolabs, Inc.) with the following conditions: 98 ^o^C (30 s), 30 cycles of 98 ^o^C (10 s), 59 ^o^C (10 s), and 72 ^o^C (10 s), and a final extension at 72 ^o^C (5 min). PCR products were run on gel (100 V for 50 min) and gel extracted and purified using the Wizard SV Gel and PCR Clean-Up System (Promega Corporation). PCR amplified and purified products of *rpoD*, *thaF*, *thaK*, and *thaQ* were sent to the RTSF Genomics Core at Michigan State University for Sanger sequencing.

RT-qPCR assays were performed using the SsoAdvanced Universal SYBR Green Supermix (Bio-Rad Laboratories, Inc.). SYBR reactions were placed into Hard-Shell PCR Plates 96-well, thin wall (Bio-Rad Laboratories, Inc. HSP9601) and analyzed using a CFX Connect Real-Time System (Bio-Rad Laboratories, Inc.). First, the dynamic range of each primer set was determined by making a 10-fold dilution series from 10 ng-0.1 pg of cDNA. The following mixture was used for each RT-qPCR assay: 10 µL SsoAdvanced universal SYBR Green supermix (2x), 0.5 µL each of forward and reverse primers, 1 µL water, and 8 µL cDNA sample (6 serially diluted samples concentrated between 1.25E-5 and 1.25 ng/uL). The RT-qPCR reaction was run with the following conditions: 95 ^o^C (3 min), 40 cycles of 95 ^o^C (10 s), 59 ^o^C (10 s), and 72 ^o^C (10 s). Following the last extension step, the melt curve was run with the following conditions: 95 ^o^C (10 s), then 65 ^o^C to 95 ^o^C in 0.5 ^o^C increments. Each primer set had a 5-fold dynamic range (10 ng-10 pg) with efficiencies between 90-110% (Table S13) The Δslope between the reference gene and each target gene were all ≤0.1, confirming that relative gene expression math models were a viable option for comparing gene expression across conditions.

cDNA concentrations across all conditions were diluted to a stock concentration of 0.0125 ng/uL. RT-qPCR reactions and conditions were prepared and run as previously described. Controls for the assay included a gDNA positive control, a no template negative control, and a no amplification (no-RT) negative control. The Livak method (2^-ΔΔC^_T_) was used to calculate relative gene expression in each test condition compared to the reference condition (*B. thailandensis*, 31 wells in M9-0.067% glucose) where target genes were *thaF*, *thaK*, and *thaQ* and the reference gene was *rpoD*.

**Supplementary Files/Datasets**

**Supplementary File 1**: Pairwise PROTEST analyses comparing the reproducibility of exometabolome profiles across biological replicate time series. Coordinates of the first two PCoA axes were used to perform PROTEST analysis in vegan. File type: .xlsx

**Supplementary File 2**: Identification of *B. thailandensis* bioactive exometabolites of interest through observation of mass spectrometry data. File type: .xlsx

**Supplementary File 3**: Genes part of the interspecies network, their corresponding modules, and GO enrichment analysis of modules. File type: .xlsx

**Supplementary File 4**: GO enrichment analysis on genes with interspecies edges from network analysis. File type: .xlsx

**Supplementary File 5**: Gene annotations for *C. subtsugae* and *P. syringae* that contained interspecies edges with *B. thailandensis* thailandamide and malleilactone biosynthetic genes, respectively. File type: .xlsx

**Supplementary File 6**: Protein alignment of the DNA starvation/stationary phase protein from *C. subtsugae* and the closest homolog in *B. thailandensis.* File type: .txt

**Supplementary File 7**: Protein alignment of the TonB-dependent siderophore receptor family protein from *P. syringae* and the closest homolog in *B. thailandensis*. File type: .txt

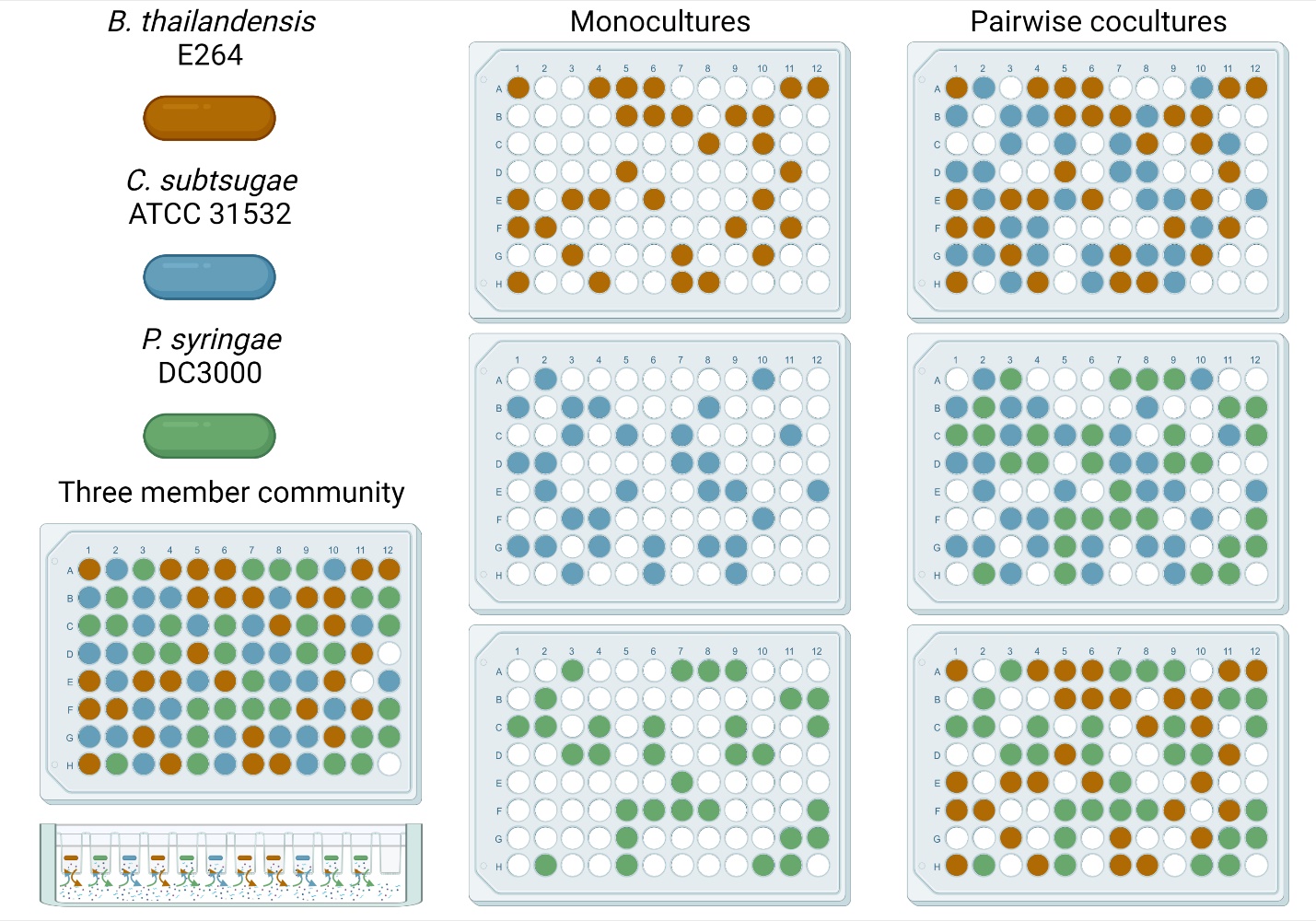

**Figure S1. Members used in synthetic community and transwell layout.** Member populations were inoculated into individual transwells. The wells contained a 0.22 µm filter that allowed for the release and transfer of exometabolites into the shared medium reservoir while maintaining physical separation of bacterial populations. There were seven possible community memberships. Each member occupied 31 wells/transwell plate regardless of community membership. Member well assignments were randomly generated for each of 4 biological replicates. Member well assignments were maintained across 1 biological replicate containing all seven possible community memberships.

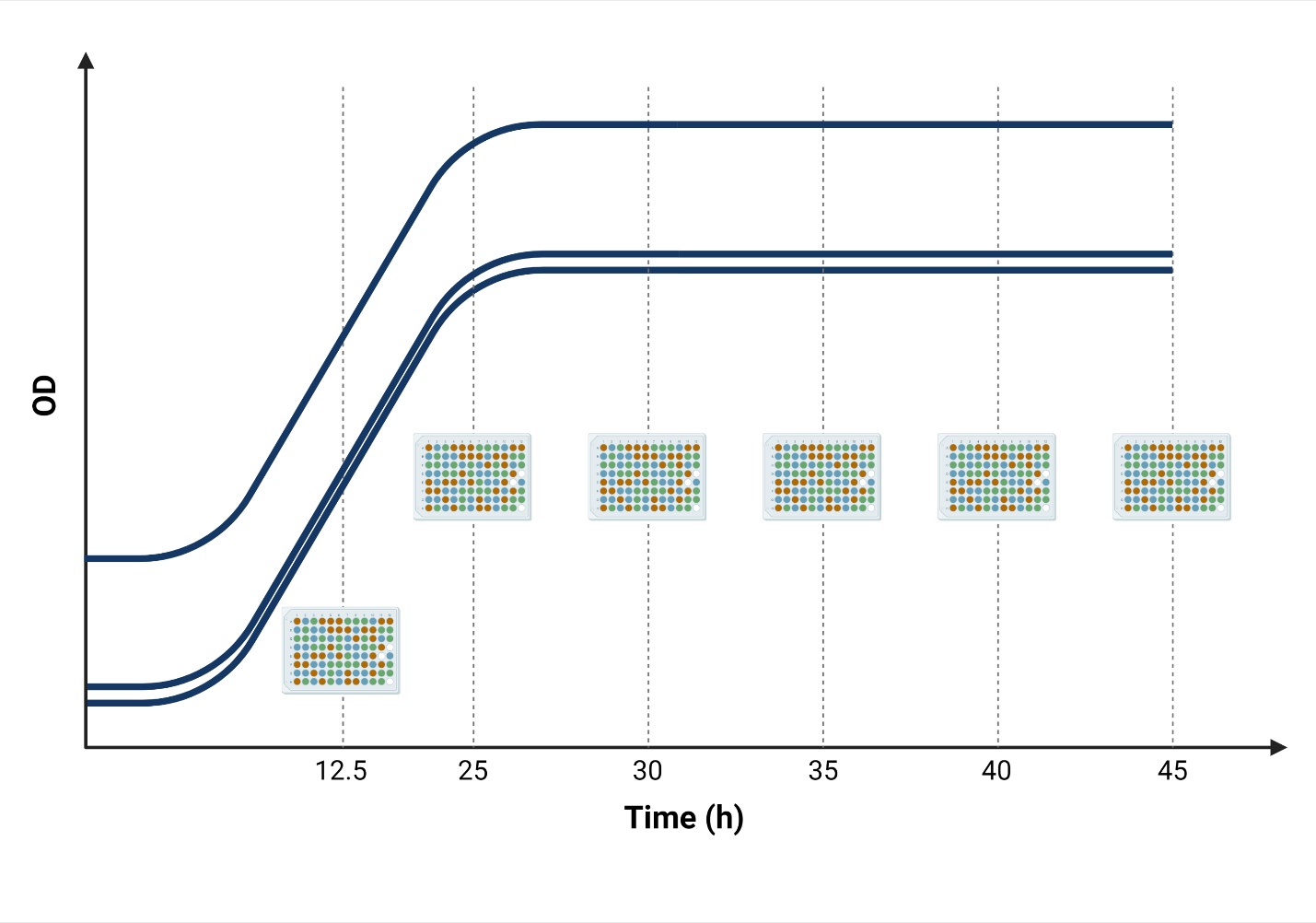
**Figure S2. Destructive sampling procedure of transwell plates.** Six replicate transwell plates were prepared for a time course experiment. At specified time points, a transwell plate was destructively sampled. Spent culture from each member was kept separate and sampled from randomized transwell wells for flow cytometry live/dead cell counts or pooled together for RNA extraction. Spent medium from the shared medium reservoir was obtained for mass spectrometry analysis. Note that all members were diluted to different starting ODs to allow for all members to achieve stationary phase within a two-hour window of each other.

**Figure S3. Differential gene expression patterns across community memberships**. Venn diagram plots of differentially expressed genes in A) *B. thailandensis* B) *C. subtsugae* and C) *P. syringae*. Differential gene expression was determined using ImpulseDE2 comparing longitudinal gene expression to a monoculture control (FDR-corrected cutoff of 0.01). Bt- *B. thailandensis*, Cs- *C.subtsugae*, and Ps – *P. syringae*.

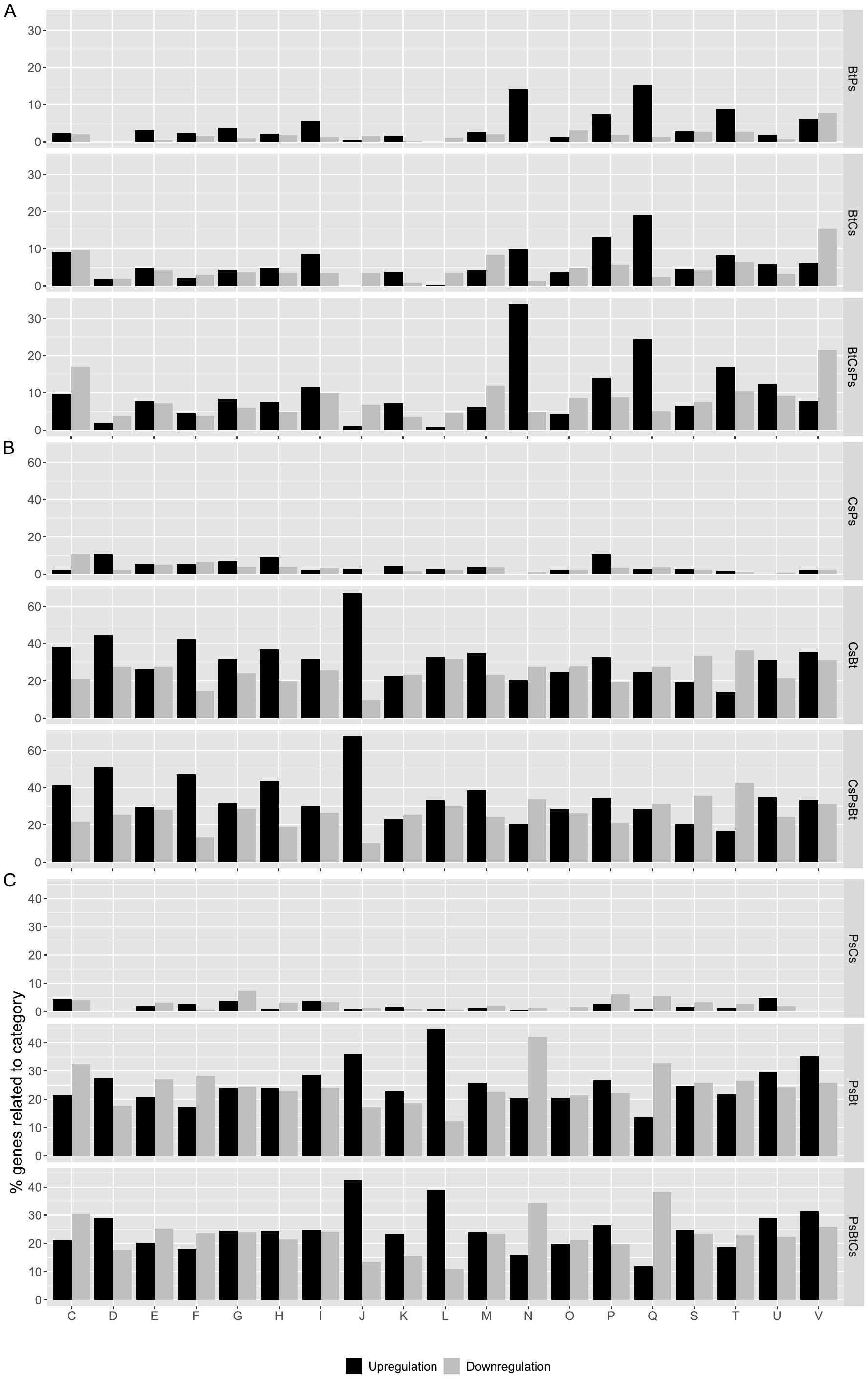

**Figure S4. Patterns of transcriptional regulation reveal biological responses to coculture**. Differentially expressed genes categorized by COG categories in A) *B. thailandensis* B) *C. subtsugae* and C) *P. syringae*. These DEGs were determined by comparing each coculture conditions to the monoculture control. COG categories include: **[C]** Energy production and conversion, **[D]** Cell cycle control, cell division, chromosome partitioning, **[E]** Amino acid transport and metabolism, **[F]** Nucleotide transport and metabolism, **[G]** Carbohydrate transport and metabolism, **[H]** Coenzyme transport and metabolism, **[I]** Lipid transport and metabolism, **[J]** Translation, ribosomal structure and biogenesis, **[K]** Transcription, **[L]** Replication, recombination and repair, **[M]** Cell wall/membrane/envelope biogenesis, **[N]** Cell motility, **[O]** Post-translational modification, protein turnover, and chaperones, **[P]** Inorganic ion transport and metabolism, **[Q]** Secondary metabolites biosynthesis, transport, and catabolism, **[S]** Function unknown, [**T]** Signal transduction mechanisms, **[U]** Intracellular trafficking, secretion, and vesicular transport, and **[V]** Defense mechanisms. Community memberships are as follows: *B. thailandensis*-*P. syringae* coculture (BtPs/PsBt), *B. thailandensis*-*C. subtsugae* coculture (BtCs/CsBt), *C. subtsugae-P. syringae* coculture (CsPs/PsCs), and the 3-member community (BtCsPs/CsPsBt/PsBtCs).

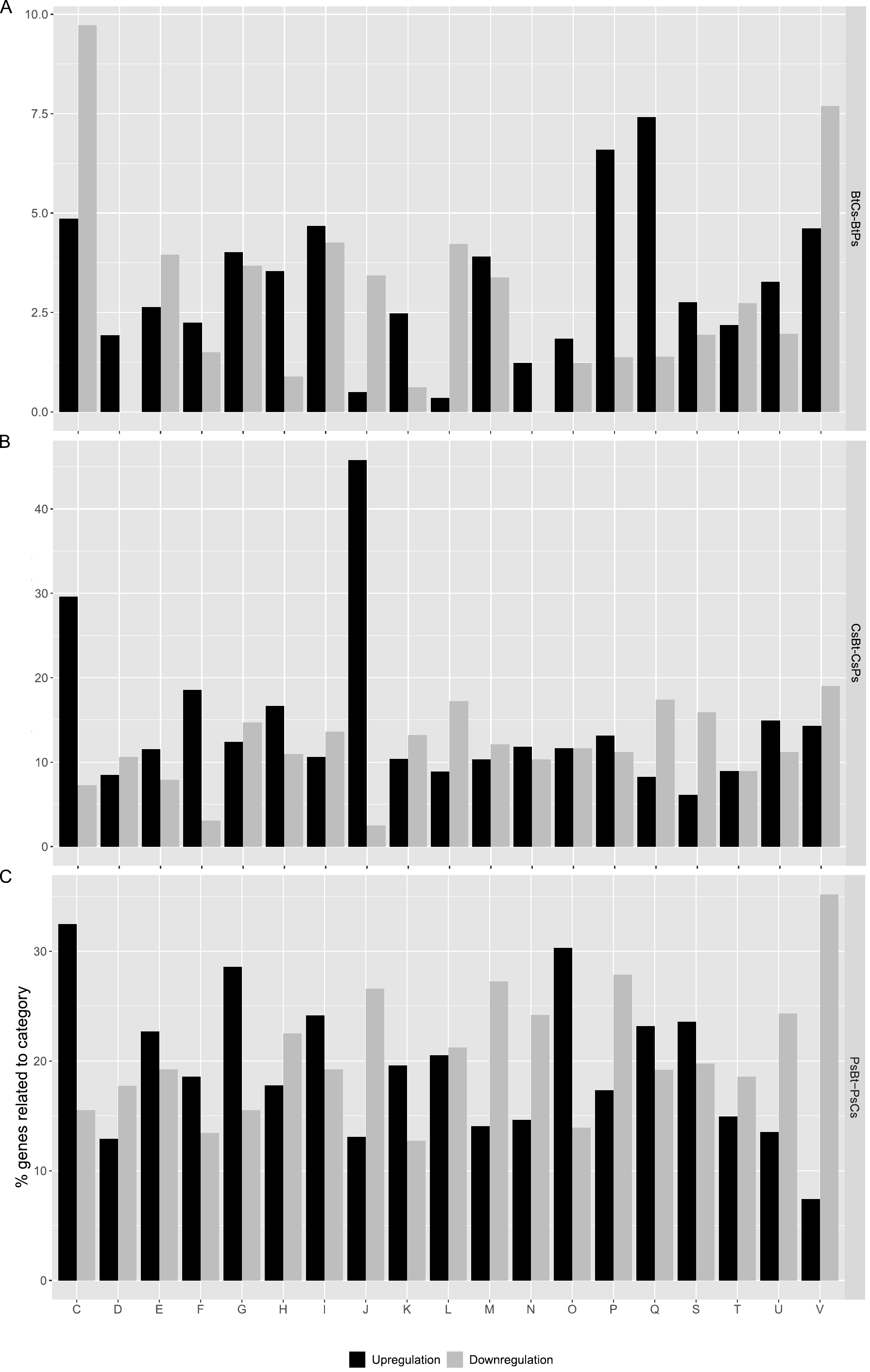

**Figure S5. Patterns of transcriptional regulation reveal pairwise coculture-specific differences**. Differentially expressed genes categorized by COG categories in A) *B. thailandensis* B) *C. subtsugae* and C) *P. syringae*. These DEGs were determined by comparing gene expression between pairwise cocultures for each member. Analyses were as follows: BtCs-BtPs; *B. thailandensis* coculture with *C. subtsugae* (case) was compared to *B. thailandensis* coculture with *P. syringae* (control), CsBt-CsPs; *C. subtsugae* coculture with *B. thailandensis* (case) was compared to *C. subtsugae* coculture with *P. syringae* (control), and PsBt-PsCs; *P. syringae* coculture with *B. thailandensis* (case) was compared to *P. syringae* coculture with *C, subtsugae* (control). COG categories are labeled in the Figure S1.4 legend.

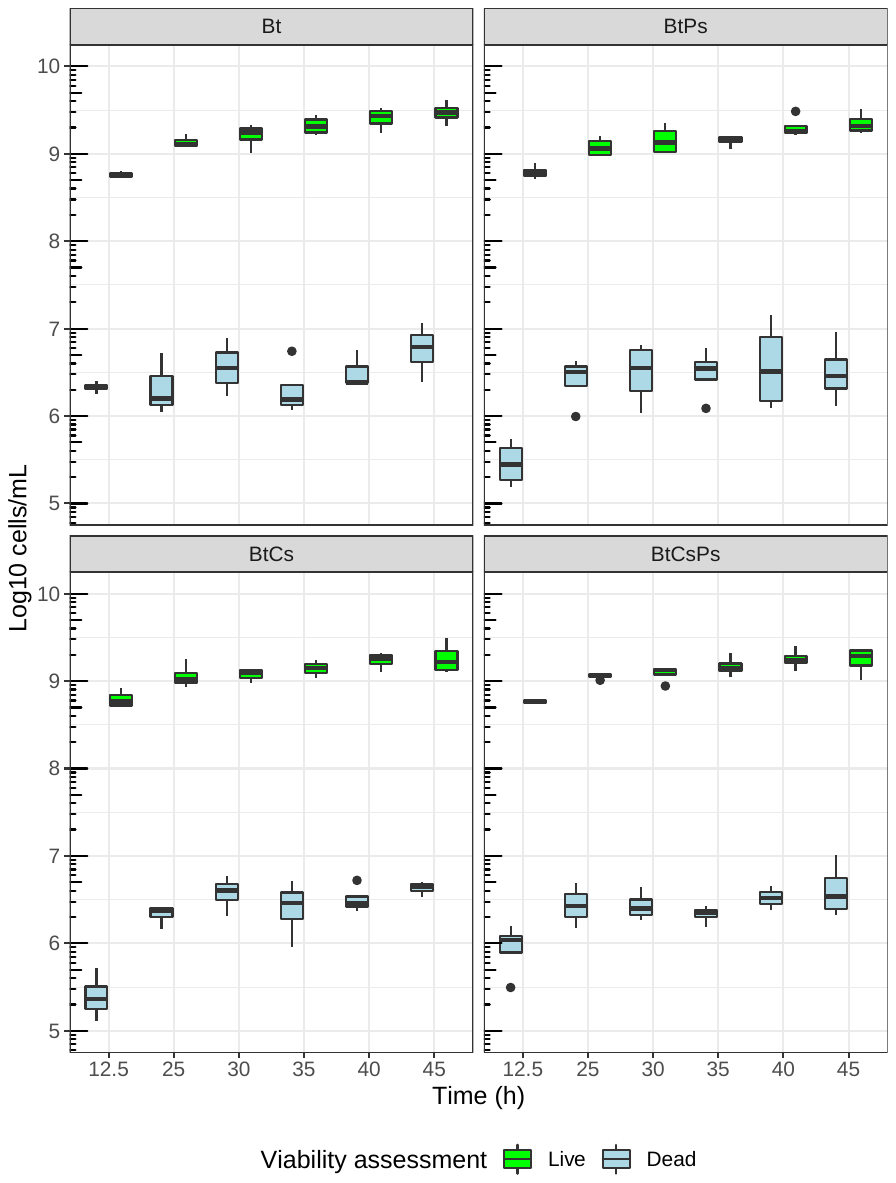

**Figure S6. *B. thailandensis* cell viability.** Counts of live (green) and dead (blue) cells throughout the time course. Cells were obtained from 5 wells in the transwell plate for 5 technical replicates/independent replicate at each time point. *B. thailandensis* monoculture (Bt), *B. thailandensis*-*P. syringae* coculture (BtPs)*, B. thailandensis*-*C. subtsugae* coculture (BtCs), and the 3-member community (BtCsPs).

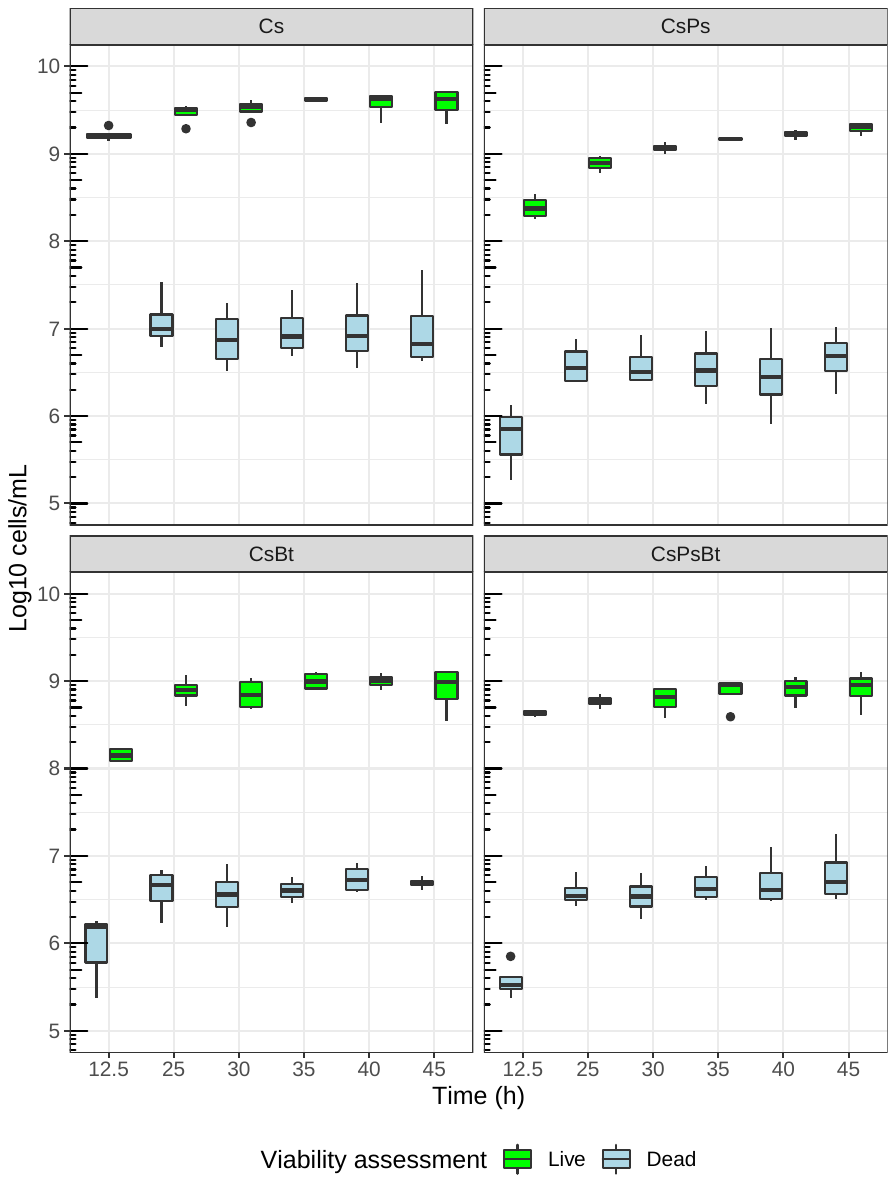

**Figure S7. *C. subtsugae* cell viability.** Counts of live (green) and dead (blue) cells throughout the time course. Cells were obtained from 5 wells in the transwell plate for 5 technical replicates/independent replicate at each time point. *C. subtsugae* monoculture (Cs), *C. subtsugae*-*P. syringae* coculture (CsPs)*, C. subtsugae*-*B. thailandensis* coculture (CsBt), and the 3-member community (CsPsBt).

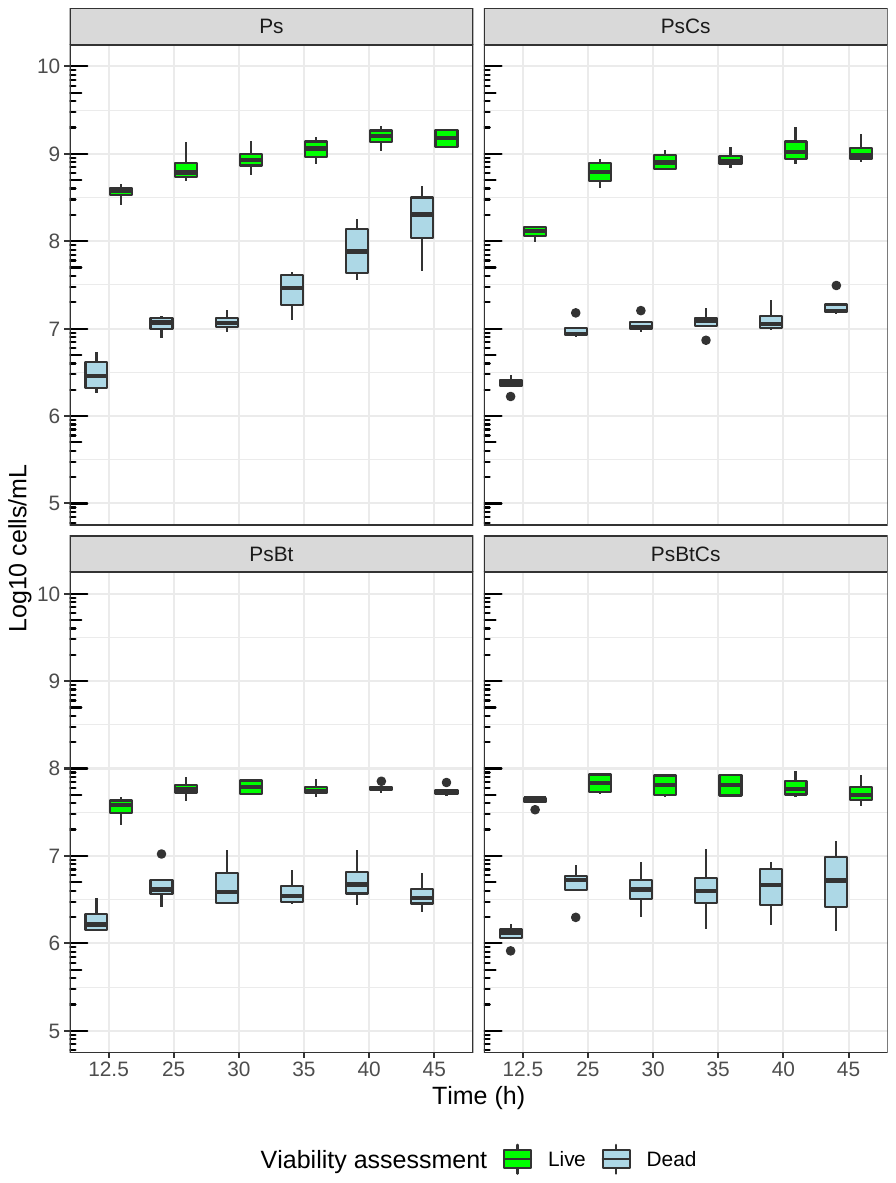

**Figure S8. *P. syringae* cell viability.** Counts of live (green) and dead (blue) cells throughout the time course. Cells were obtained from 5 wells in the transwell plate for 5 technical replicates/independent replicate at each time point. *P. syringae* monoculture (Ps), *P. syringae***-***C. subtsugae* coculture (PsCs)*, P. syringae*-*B. thailandensis* coculture (PsBt), and the 3-member community (PsBtCs).

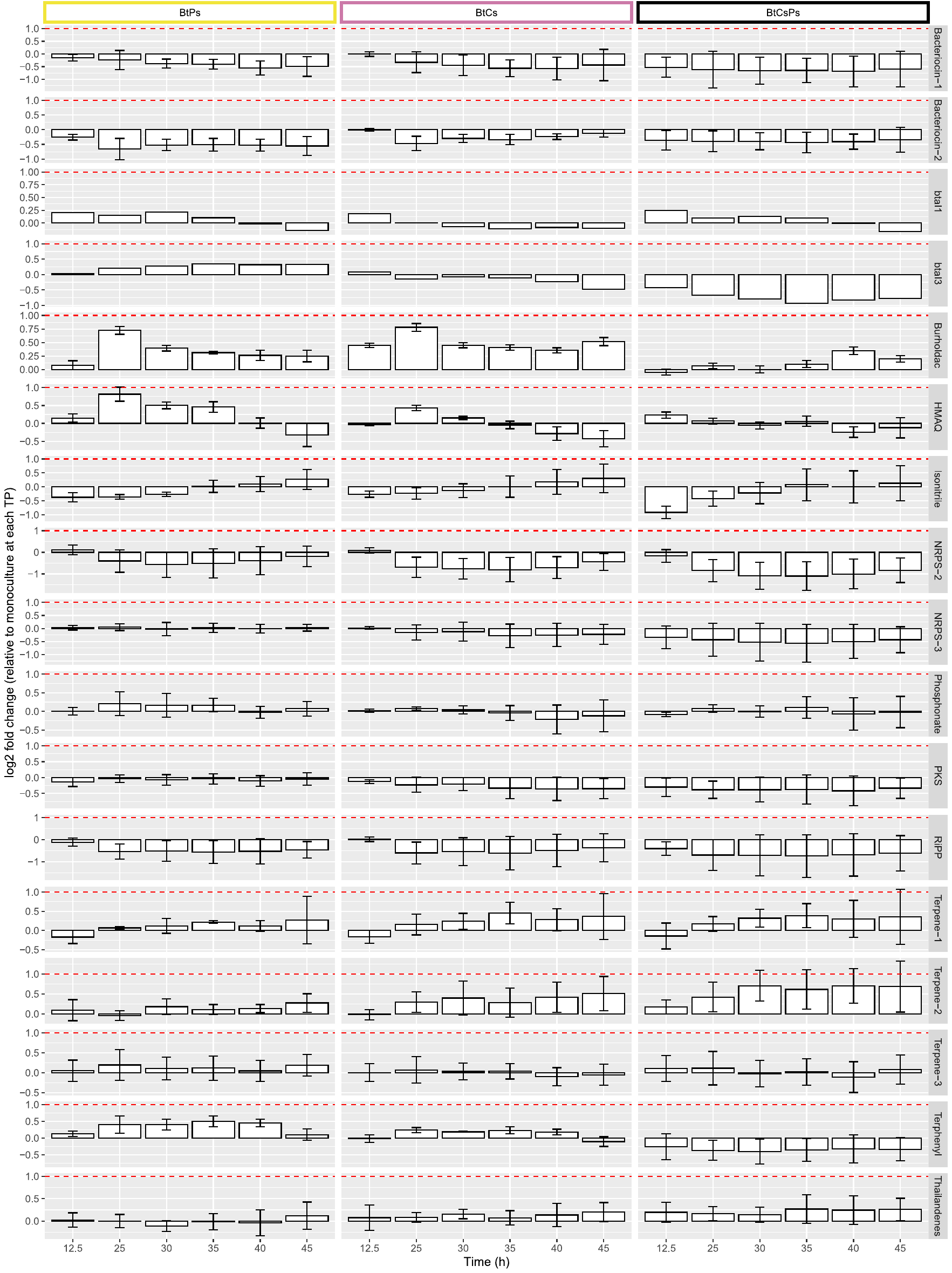

**Figure S9. BSGC downregulated or unaltered in *B. thailandensis*.** Biosynthetic genes involved in each BSGC were determined with antiSMASH and evidence from literature. At each timepoint, the average log2 fold-change (LFC) was determined across all biosynthetic genes for each BSGC. The horizontal line represents a LFC threshold of 1. Note that plots for each BSGC have separate scales for the Y-axis.

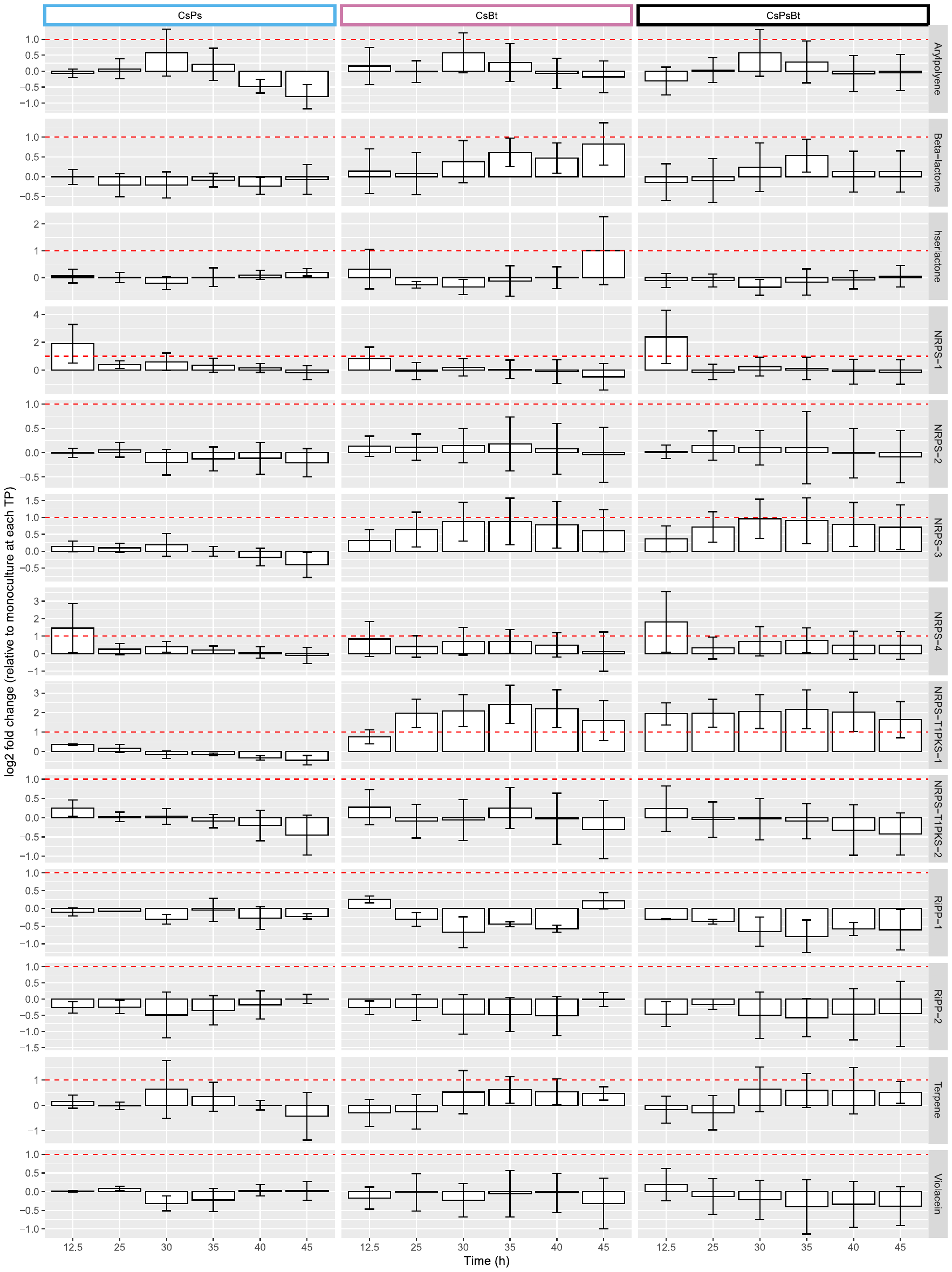

**Figure S10. Patterns of transcriptional regulation for BSGC in *C. subtsugae*.** Biosynthetic genes involved in each BSGC were determined with antiSMASH and evidence from literature. At each timepoint, the average log2 fold-change (LFC) was determined across all biosynthetic genes for each BSGC. The horizontal line represents a LFC threshold of 1. Note that plots for each BSGC have separate scales for the Y-axis.

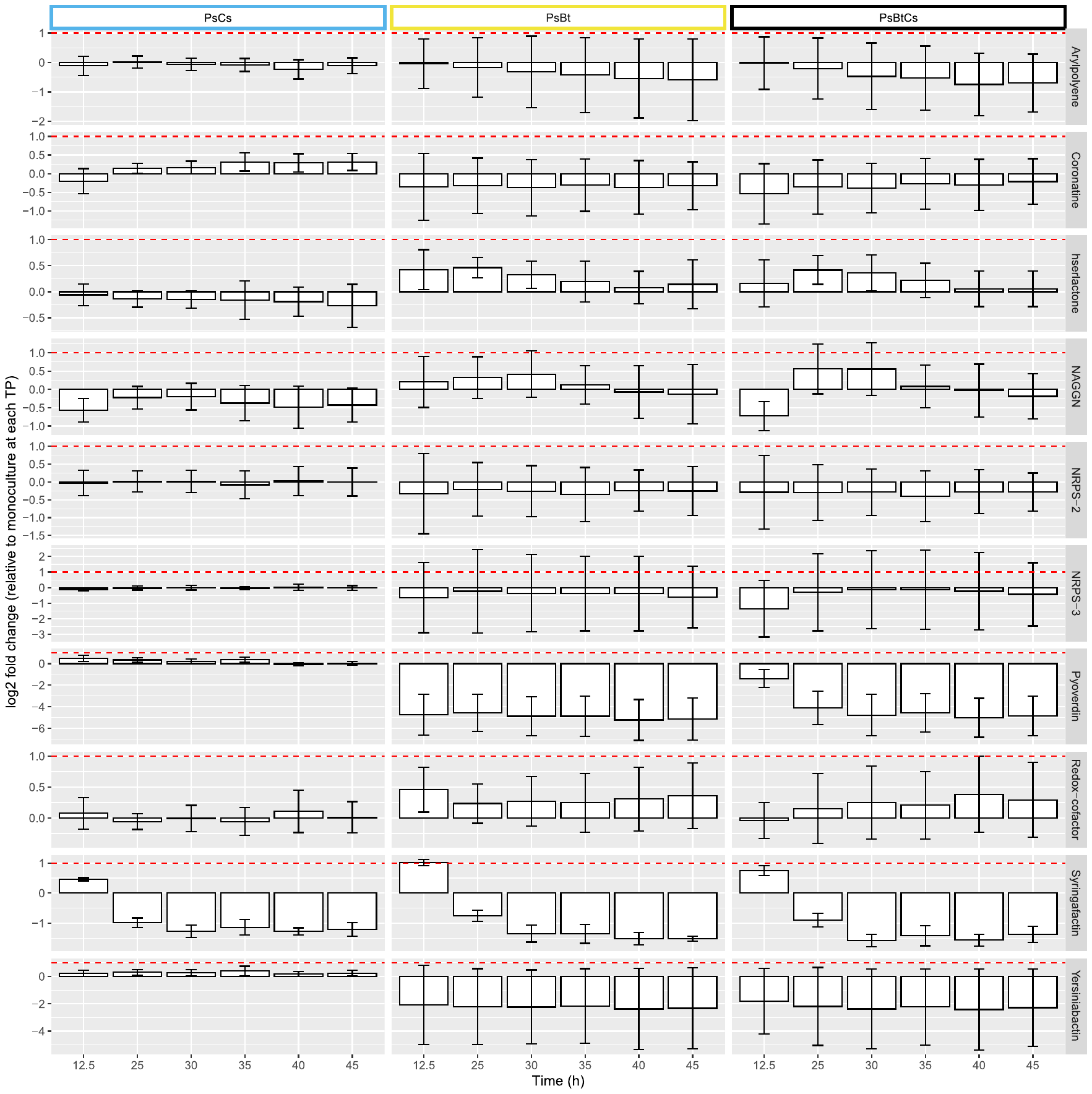

**Figure S11. Patterns of transcriptional regulation for BSGC in *P. syringae*.** Biosynthetic genes involved in each BSGC were determined with antiSMASH and evidence from literature. At each timepoint, the average log2 fold-change (LFC) was determined across all biosynthetic genes for each BSGC. The horizontal line represents a LFC threshold of 1. Note that plots for each BSGC have separate scales for the Y-axis.

**Figure S12. Flow diagram for interspecies co-expression network analysis.** An interspecies coexpression network was created based on transcript counts from *B. thailandensis*-*C. subtsugae* and *B. thailandensis*-*P. syringae* cocultures. All genes that passed initial quality filtering were included in the analysis to generate networks. Unweighted gene coexpression networks were generated with a Z-score cutoff of 4.5. Intraspecies genes were used to identify network modules. Gene ontology enrichment analysis was performed on nodes with interspecies edges.

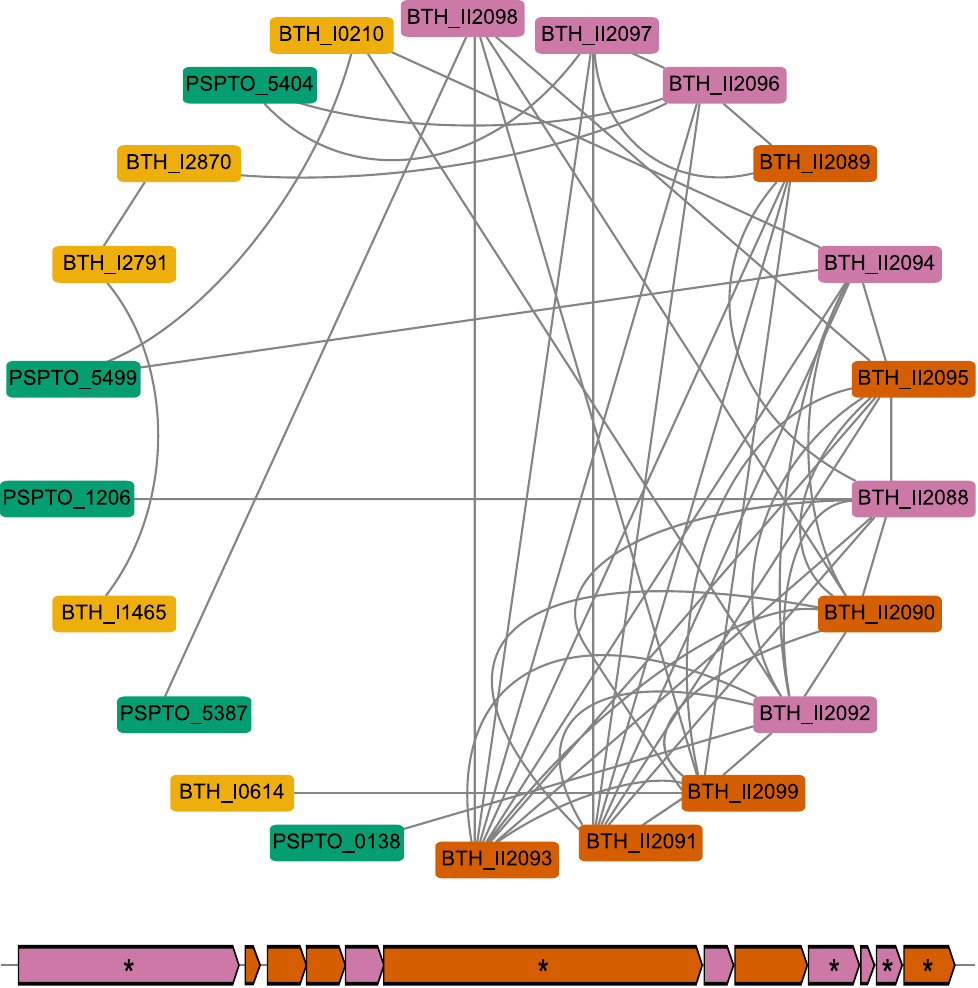

**Figure S13. *B. thailandensis* genes involved in malleilactone production are detected as interspecies edges in the *B. thailandensis*-*P. syringae* coexpression network and biosynthetic genes organize into network modules.** A network module containing the malleilactone BSGC is shown. The network module nodes are color coded by *B.* *thailandensis* gene type (BSGC or not) and type of connections (interspecies or not): malleilactone biosynthetic genes that had interspecies edges (magenta), malleilactone biosynthetic genes that did not have interspecies edges (orange), or other genes that were not part of the BSGC (yellow); as well as genes that were from *P. syringae* (green). The chromosomal organization of the malleilactone BSGC is shown below the network module. The same colors are applied to the BSGC operons. Asterisks indicate core biosynthetic genes in the BSGCs, as predicted from antiSMASH.

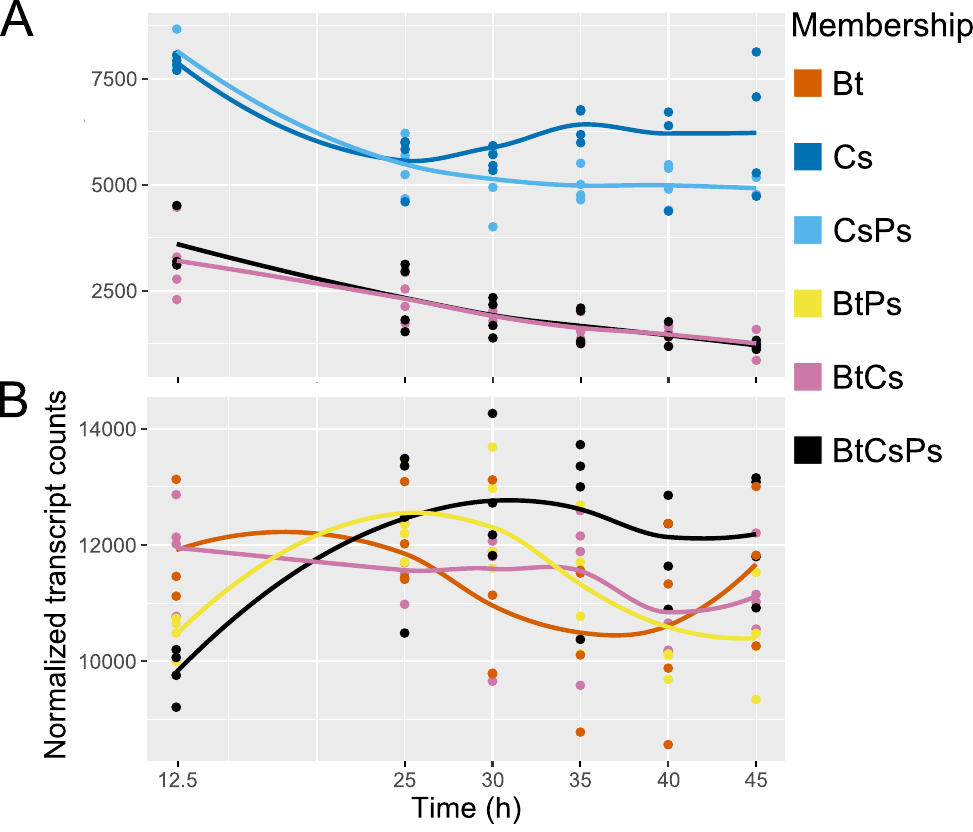

**Figure S14. The DNA starvation/stationary phase protection gene, *dpsA*, in downregulated in *C. subtsugae* when cocultured with *B. thailandensis* while unaltered in *B. thailandensis*.** Transcript abundance trajectories of *dpsA* are plotted for *C. subtsugae* (A) and *B. thailandensis* (B). Time course scatter plots were smooth curve fitted by loess. Community memberships are as follows: *B. thailandensis* monoculture (Bt), *C. subtsugae* monoculture (Cs), *C. subtsugae-P. syringae* coculture (CsPs), *B. thailandensis-P. syringae* coculture (BtPs), *B. thailandensis-C. subtsugae* coculture (BtCs), and the 3-member community (BtCsPs).

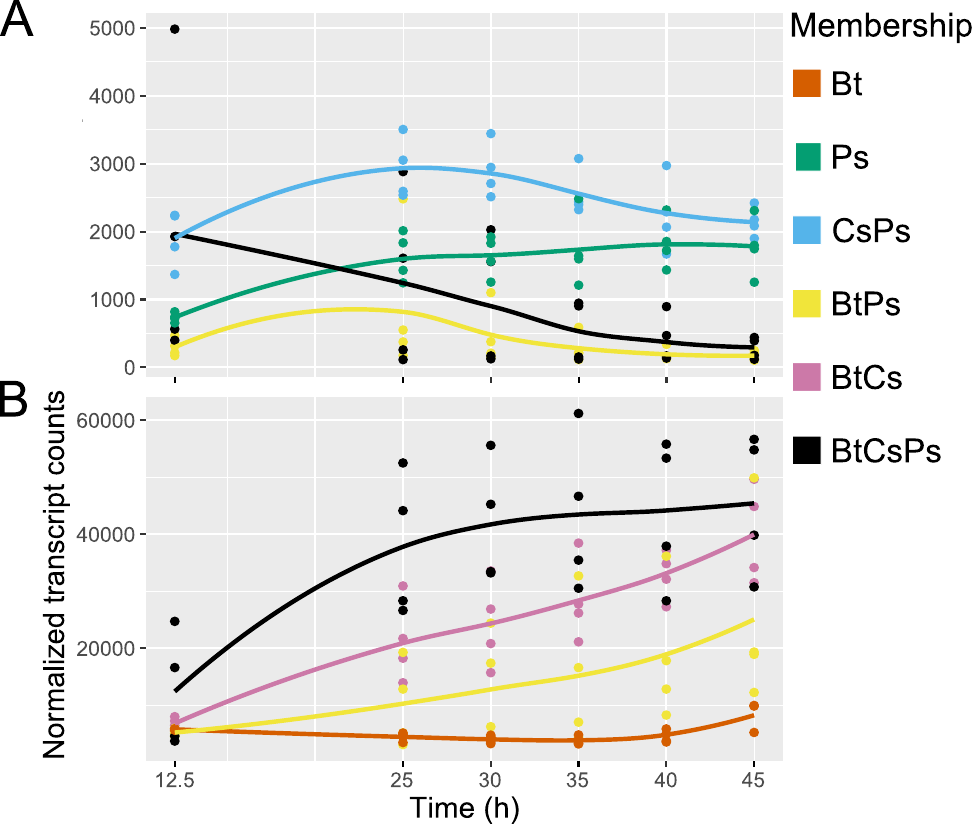

**Figure S15. The gene encoding a TonB-dependent siderophore receptor is downregulated in *P. syringae* when cocultured with *B. thailandensis* while upregulated in *B. thailandensis*.** Transcript abundance trajectories of *tonB* are plotted for *P. syringae* (A) and *B. thailandensis* (B). Time course scatter plots were smooth curve fitted by loess. Community memberships are as follows: *B. thailandensis* monoculture (Bt), *P. syringae* monoculture (Ps), *C. subtsugae-P. syringae* coculture (CsPs), *B. thailandensis-P. syringae* coculture (BtPs), *B. thailandensis-C. subtsugae* coculture (BtCs), and the 3-member community (BtCsPs).

**Supplementary Table S1.** Percent variation explained on the effect of membership, time, and their interaction on transcriptomic profiles.

|  | Membership | Time | Membership x Time |
| --- | --- | --- | --- |
| *B. thailandensis* | 46.26 | 13.24 | 63.11 |
| C. *subtsugae* | 60.60 | 3.88 | 68.29 |
| *P. syringae* | 77.03 | 0.00 | 81.40 |

**Supplementary Table S2.** Summary of Protest analyses comparing transcriptional profiles through time across independent replicates. Coordinates of the first two PCA axes were used to perform Protest analyses. Ranges reflect separate Protest analyses performed between all replicates in a community membership. Values in parenthesis represent the median *P* value.

|  | m12 | R | *P* |
| --- | --- | --- | --- |
| *B. thailandensis* |  |  |  |
| Monoculture | 0.048 – 0.820 | 0.424 – 0.976 | 0.010 – 0.867 (0.200) |
| *P. syringae* coculture | 0.018 – 0.049 | 0.975 – 0.991 | 0.001 – 0.001 (0.001) |
| *C. subtsugae* coculture | 0.010 – 0.112 | 0.943 – 0.995 | 0.001 – 0.003 (0.001) |
| 3-member | 0.013 – 0.162 | 0.916 – 0.994 | 0.001 – 0.006 (0.001) |
| *C. subtsugae* |  |  |  |
| Monoculture | 0.011 – 0.140 | 0.927 – 0.995 | 0.004 – 0.067 (0.039) |
| *P. syringae* coculture | 0.045 – 0.206 | 0.891 – 0.977 | 0.003 – 0.042 (0.008) |
| *B. thailandensis* coculture | 0.091 – 0.182 | 0.905 – 0.954 | 0.001 – 0.108 (0.019) |
| 3-member | 0.190 – 0.543 | 0.676 – 0.900 | 0.001 – 0.208 (0.013) |
| *P. syringae* |  |  |  |
| Monoculture | 0.178 – 0.538 | 0.680 – 0.907 | 0.008 – 0.136 (0.054) |
| *C. subtsugae* coculture | 0.035 – 0.251 | 0.865 – 0.982 | 0.001 – 0.083 (0.021) |
| *B. thailandensis* coculture | 0.021 – 0.290 | 0.843 – 0.990 | 0.001 – 0.001 (0.001) |
| 3-member | 0.034 – 0.687 | 0.560 – 0.983 | 0.007 – 0.317 (0.038) |

**Supplementary Table S3.** PERMANOVA results calculated on independently replicated time series within members across all community memberships. PERMANOVA results are presented as *P* values, R^2^ values, and pseudo-F statistic results in the first row. Post-hoc pairwise PERMANOVA results are presented below the first row.

|  | *B. thailandensis* | *C. subtsugae* | *P. syringae* | |
| --- | --- | --- | --- | --- |
| adonis | *P* = 0.001,  R^2^ = 0.480,  F = 27.686 | *P* = 0.001,  R^2^ = 0.619,  F= 47.15 | | *P* = 0.001,  R^2^ = 0.778,  F= 107.21 |
| Monoculture vs  *B. thailandensis* coculture | - | 0.002 | | 0.001 |
| Monoculture vs  *C. subtsugae* coculture | 0.001 | - | | 0.001 |
| Monoculture vs  *P. syringae* coculture | 0.001 | 0.010 | | - |
| Monoculture vs  3-member | 0.001 | 0.002 | | 0.001 |
| *B. thailandensis* coculture vs  *C. subtsugae* coculture | - | - | | 0.001 |
| *B. thailandensis* coculture vs  *P. syringae* coculture | - | 0.002 | | - |
| *C. subtsugae* coculture vs  *P. syringae* coculture | 0.001 | - | | - |
| *B. thailandensis* coculture vs  3-member | - | 0.248 | | 0.068 |
| *C. subtsugae* coculture vs  3-member | 0.001 | - | | 0.001 |
| *P. syringae* coculture vs  3-member | 0.001 | 0.002 | | - |

**Supplementary Table S4.** Percent variation explained on the effect of membership, time, and their interaction on exometabolite profiles.

|  | Membership | Time | Membership x Time |
| --- | --- | --- | --- |
| Polar Positive | 45.76 | 7.26 | 55.89 |
| Polar Negative | 51.61 | 4.12 | 58.83 |
| Nonpolar Positive | 56.92 | 9.49 | 71.88 |
| Nonpolar Negative | 64.77 | 7.94 | 79.38 |

**Supplementary Table S5.** Summary of Protest analyses comparing exometabolite composition through time across independent replicates. Coordinates of the first two PCoA axes were used to perform Protest analyses. Ranges reflect separate Protest analyses performed for each polarity (polar/nonpolar) and ionization mode (positive/negative).

|  | m12 | R | *P* |
| --- | --- | --- | --- |
| *C. subtsugae*-*P. syringae* coculture | 0.022 – 0.906 | 0.307 – 0.989 | 0.001 – 0.849 (0.025) |
| *B. thailandensis*-*P. syringae* coculture | 0.015 – 0.592 | 0.638 – 0.992 | 0.001 – 0.667 (0.050) |
| *B. thailandensis-C. subtsugae* coculture | 0.003 – 0.456 | 0.738 – 0.995 | 0.001 – 0.250 (0.003) |
| 3-member community | 0.021 – 0.556 | 0.667 – 0.990 | 0.001 – 0.133 (0.003) |

**Supplementary Table 6.** PERMANOVA results calculated on independently replicated time series across coculture community memberships. PERMANOVA results are presented as *P* values, R^2^ values, and pseudo-F statistic results in the first row. Post-hoc pairwise PERMANOVA results are presented below the first row.

|  | Polar Positive | Polar Negative | Nonpolar Positive | Nonpolar Negative |
| --- | --- | --- | --- | --- |
| adonis | *P* = 0.001, R^2^ = 0.475, F = 27.711 | *P* = 0.001, R^2^ = 0.531, F = 34.773 | *P* = 0.001 R^2^ = 0.585, F = 37.549 | *P* = 0.001, R^2^ = 0.662, F = 45.743 |
| *B. thailandensis*-*C. subtsugae* coculture vs  *C. subtsugae*-*P. syringae* coculture | 0.001 | 0.001 | 0.001 | 0.001 |
| *B. thailandensis*-*C. subtsugae* coculture vs  *B. thailandensis*- *P. syringae* coculture | 0.001 | 0.001 | 0.001 | 0.001 |
| *B. thailandensis*- *P. syringae* coculture vs  *C. subtsugae*-*P. syringae* coculture | 0.001 | 0.001 | 0.001 | 0.001 |
| 3-member community vs  *C. subtsugae*-*P. syringae* coculture | 0.001 | 0.001 | 0.001 | 0.001 |
| 3-member community vs  *B. thailandensis*- *P. syringae* coculture | 0.001 | 0.001 | 0.001 | 0.001 |
| 3-member community vs  *B. thailandensis*-*C. subtsugae* coculture | 0.002 | 0.008 | 0.019 | 0.025 |

**Supplementary Table S7.** Number of predicted biosynthetic gene clusters (BSGCs, first row) followed by the quantity of upregulated BSGCs in cocultures.

|  | *B. thailandensis* | *C. subtsugae* | *P. syringae* |
| --- | --- | --- | --- |
| Predicted BSGCs | 28 | 14 | 10 |
| *C. subtsugae*-*P. syringae* coculture | - | 0 | 0 |
| *B. thailandensis*-*P. syringae* coculture | 8 | - | 0 |
| *B. thailandensis*-*C.* *subtsugae* coculture | 10 | 1 | - |
| 3-member community | 11 | 1 | 0 |

**Supplementary Table S8.** One-way ANOVA^a^ comparing the quantitation of identified secondary metabolites between community memberships with *B. thailandensis* membership.

|  | Df (between) | Df (within) | *F* value | *p* |
| --- | --- | --- | --- | --- |
| Bactobolin | 6 | 160 | 392.10 | <2e-16 |
| Capistruin | 6 | 160 | 77.83 | <2e-16 |
| Melleilactone | 6 | 121 | 150.10 | <2e-16 |
| Rhamnolipid^b^ | 6 | 136 | 39.34 | <2e-16 |
| Thailandamide | 6 | 121 | 61.02 | <2e-16 |
| Pyochelin | 6 | 136 | 105.20 | <2e-16 |

^a^Formula: aov(formula = log(Value) ~ Membership, data = .)

^b^Rhamnolipid congener Rha-Rha-C14-C14

**Supplementary Table S9.** TukeyHSD post-hoc results comparing quantitation of identified secondary metabolites between community memberships with *B. thailandensis* membership. Values represent the adjusted P-value.

|  | Bactobolin | Capistruin | Melleilactone | Rhamnolipid^a^ | Thailandamide | Pyochelin |
| --- | --- | --- | --- | --- | --- | --- |
| Monoculture vs *P. syringae* coculture | 8.41E-01 | 8.00E-07 | 4.00E-07 | 2.26E-01 | 1.25E-04 | 4.85E-01 |
| Monoculture vs *C. subtsugae* coculture | 3.54E-01 | 8.45E-02 | < 1.00E-07 | 2.53E-02 | < 1.00E-07 | 8.55E-01 |
| Monoculture vs 3-member | 8.27E-01 | 5.00E-07 | < 1.00E-07 | 1.00E-05 | < 1.00E-07 | 7.13E-04 |
| *C. subtsugae* coculture vs *P. syringae* coculture | 8.38E-01 | 6.48E-03 | 5.45E-02 | 7.68E-01 | 9.27E-03 | 9.11E-01 |
| *P. syringae* coculture vs 3-member | 3.28E-01 | 9.99E-01 | 1.00E-07 | 6.50E-03 | 4.32E-03 | 4.24E-02 |
| *C. subtsugae* coculture vs 3-member | 6.08E-02 | 4.20E-03 | 1.76E-03 | 7.76E-02 | 9.97E-1 | 6.20E-03 |

^a^Rhamnolipid congener Rha-Rha-C14-C14

**Supplementary Table S10.** Relative gene expression of genes in the thailandamide operon across different SynCom conditions. *B. thailandensis* (31 wells) in M9-0.067% glucose was the control condition and *rpoD* was the reference gene.

|  | *thaF* | *thaK* | *thaQ* |
| --- | --- | --- | --- |
| Bt (62 wells) | 0.523 | 0.550 | 0.650 |
| Bt (93 wells) | 0.303 | 0.138 | 0.188 |
| Bt-Ps (31 wells/member) | 1.311 | 1.675 | 2.163 |
| Bt-Cv (31 wells/member) | 2.375 | 2.304 | 1.05 |
| Bt-Cv-Ps (31 wells/member) | 2.048 | 2.433 | 1.742 |

**Supplementary Table S11.** Network summary results from interspecies coexpression networks.

| Network | *B. thailandensis-C. subtsugae* | | *B. thailandensis-P. syringae* | |
| --- | --- | --- | --- | --- |
| Member | *B. thailandensis* | *C. subtsugae* | *B. thailandensis* | *P. syringae* |
| Total nodes | 2701 | 2043 | 3254 | 3478 |
| Nodes with only intraspecies edges | 2418 | 1814 | 2749 | 2996 |
| Nodes with interspecies edges | 283 | 229 | 505 | 482 |
| Total edges | 9382 | 7240 | 15801 | 23319 |
| Intraspecies edges | 9074 | 6932 | 15056 | 22574 |
| Interspecies edges | 308 | | 745 | |

**Supplementary Table S12.** Primers used for RT-qPCR analysis of genes in the thailandamide operon.

| Primer | Sequence (5’ > 3’) | Product size (bps) | Reference |
| --- | --- | --- | --- |
| *rpoD*_F | ACCGTCGTGGCTACAAATTC | 117 | [3] |
| *rpoD*_R | TCGTCTCGATCATGTGAACC |  |  |
| *thaF*_F | CATGCACGCGTTTCTGTTTC | 113 | This study |
| *thaF*_F | TCGTAGCCCAAGATCTCGTT |  |  |
| *thaK*_F | GGTATTGAGGCCATGAACGT | 104 | This study |
| *thaK*_F | CATCAGCAGATTCGCGAAAC |  |  |
| *thaQ*_F | GAACGCGTCGAAGGATTTTC | 115 | This study |
| *thaQ*_F | ATTCGTTCGGGTACTTCTGC |  |  |

**Supplementary Table S13.** RT-qPCR efficiencies of reference and target genes for relative expression analysis of genes in the thailandamide operon.

| Gene | Slope | R^2^ | Efficiency (%) |
| --- | --- | --- | --- |
| *rpoB* | -3.424 | 0.999 | 102.6 |
| *thaF* | -3.204 | 0.996 | 105.2 |
| *thaK* | -3.358 | 0.995 | 98.5 |
| *thaQ* | -3.305 | 0.997 | 100.7 |
